# Allosteric cooperation in ß-lactam binding to a non-classical transpeptidase

**DOI:** 10.1101/2021.09.06.459080

**Authors:** Nazia Ahmad, Sangita Kachhap, Varsha Chauhan, Kunal Sharma, Pallavi Juneja, C. Korin Bullen, Tomasz Borowski, William R. Bishai, Gyanu Lamichhane, Pankaj Kumar

## Abstract

*Mycobacterium tuberculosis* peptidoglycan (PG) is atypical as its synthesis involves a new enzyme class, L,D-transpeptidases. Prior studies of L,D-transpeptidases have identified only the catalytic site that binds to peptide moiety of the PG substrate or ß-lactam antibiotics. This insight was leveraged to develop mechanism of its activity and inhibition by ß-lactams. Here we report identification of an allosteric site at a distance of 21 Å from the catalytic site that binds the sugar moiety of PG substrates (hereafter referred to as the S-pocket). This site also binds a second ß-lactam molecule and influences binding at the catalytic site. We provide evidence that two ß-lactam molecules bind co-operatively to this enzyme, one non-covalently at the S-site and one covalently at the catalytic site. This dual ß-lactam binding phenomenon is previously unknown and is an observation that may offer novel approaches for the structure-based design of new ß-lactam antibiotics for *M. tuberculosis.*

## INTRODUCTION

Tuberculosis (TB), is a major threat to global health as it claims more human lives than any other bacterial infection (Chakaya et al., 2021). The emergence of multi-(MDR) and extensively drug-resistant (XDR) strains of *Mycobacterium tuberculosis (M.tb)*, the bacterium that causes TB, has further limited our capability to fight the disease. One major factor contributing to the emergence of drug-resistant TB is poor compliance to prolonged treatment regimens. While combinatorial drug therapy kills the majority of *M.tb* bacilli a subset of the bacterial population, defined as “persisters”, tolerate TB drugs. This persister subset requires prolonged treatment for sterilization to occur (Gideon and Flynn, 2011; Lillebaek et al., 2002; Peddireddy et al., 2017; Wayne and Hayes, 1996). Mechanisms of persistence in *M.tb are* likely multifactorial, involving cell wall peptidoglycan remodelling, transporters or efflux pumps, and alternative energy sources (Keren et al., 2011; Zhang et al., 2012). Molecular understanding of these pathways may facilitate discovery of new therapeutics to overcome the current challenges posed by *M.tb* persistence and drug-resistance.

Peptidoglycan (PG) is an essential component of the bacterial cell wall and constitutes the exoskeleton of bacterial cells. PG consist of long glycan chains composed of two different sugars N-acetyl muramic acid (NAM) and N-acetyl glucose amine (NAG) that are crossed linked via short stem peptide chains. The PG composition of *M.tb* in slowly-replicating states is likely to be distinct from the one during active growth (Gupta et al., 2010; Schoonmaker et al., 2014; Wietzerbin et al., 1974). In particular, a high percentage of peptide cross-links in *M.tb* join the third amino acids (3-3 linkages) of the adjacent stem peptides instead of the classical 4-3 linkages, and these linkages are formed by transpeptidases (Tolufashe et al., 2020). The 4-3 linkages, which were historically considered to predominate throughout bacterial growth and senescence, are generated by a well-known enzyme class, namely the D,D-transpeptidases (also known as penicillin-binding proteins) (Tolufashe et al., 2020). The 3-3 linkages are generated by the more recently discovered enzyme class, the L,D-transpeptidases (Mainardi et al., 2005).

Among the five L,D-transpeptidase paralogs of *M.tb*, Ldt_Mt2_ plays an important role since an *M.tb* strain lacking the gene encoding this enzyme exhibits attenuation of persistence and virulence (Bianchet et al., 2017; Brammer Basta et al., 2015; Dubee et al., 2012; Gupta et al., 2010; Libreros-Zuniga et al., 2019; Sanders et al., 2014; Schoonmaker et al., 2014). Additional reports have demonstrated altered and attenuated cellular physiology of *M.tb* in association with the loss of function of LdtMt1 (Schoonmaker et al., 2014) and LdtMt5(Brammer Basta et al., 2015). The necessity of L,D-transpeptidases for virulence has suggested that they may comprise valuable drug targets, and indeed these enzymes are inhibited by the carbapenem class of ß-lactam drugs (Mainardi et al., 2005). Recent work further suggests the efficacy of these carbapenems against both dividing and non-dividing mycobacteria (Hugonnet et al., 2009). To further evaluate the druggability of the L,D-transpeptidases class, several independent groups have described the crystal structures of this enzyme class bound to carbapenems such as meropenem, tebipenem, biapenem and faropenem (Bianchet et al., 2017; Erdemli et al., 2012; Kim et al., 2013; Li et al., 2013; Steiner et al., 2017). The biochemical mechanisms and kinetics of inhibition of L,D-transpeptidases by ß-lactams have been documented (Cordillot et al., 2013), and new experimental ß-lactams that target *M.tb* L,D-transpeptidases have recently been described (Bianchet et al., 2017; Kumar et al., 2017; Martelli et al., 2021).

The L,D-transpeptidase class in *M.tb* are comprised of at least four substructural domains including two immunoglobulin-like domains (IgD1 & IgD2), a YkuD domain, and a C-terminal subdomain (CTSD). The YkuD domain is known to play a role in catalytic function and ß-lactam binding (Erdemli et al., 2012), while the roles of the other domains remains less certain. The YkuD domain has a highly conserved motif HXX14-17[S/T]HGChN containing three residues analogous to the catalytic triad of cysteine proteases: a cysteine, a histidine and a third residue (Cystine 354, Histidine 336 and Serine 337 in Ldt_Mt2_) to catalyse the transpeptidation reaction (Erdemli et al., 2012). This catalytic triad residues resides under a flap formed by a long loop. The flap can open and close to create two cavities (the inner and outer cavities), around a cysteine residue, that are connected by a narrow tunnel (Fakhar et al., 2017). It is proposed that these cavities are binding sites for the acyl acceptor and acyl donor tetrapeptide stems (L-Alanyl-D-Glutamyl-*meso*-diaminopimelyl-D-alanine) with the donor tetrapeptide binding to the outer cavity and the acceptor tetrapeptide to the inner cavity (Erdemli et al., 2012). As the ß-lactam class of drugs mimics the tetrapeptide stems of PG, several of the carbapenems drugs have been found to bind both the inner and outer cavities to form covalent linkage with catalytic cysteine residue of the L,D-transpeptidase (Bianchet et al., 2017; Kim et al., 2013; Kumar et al., 2017).

Despite the significance of L,D-transpeptidases in *M.tb* cell physiology and TB disease, the structural and molecular details of how different chemical groups of the nascent PG structure interact with this enzyme class are not sufficiently understood. The disaccharide-tetrapeptide *N-*acetylglucosamine-*N-*acetylmuramic acid-L-Alanyl-D-Glutamyl-*meso*-diaminopimelyl-D-alanine is the substrate for L,D-transpeptidases (Cordillot et al., 2013; Lavollay et al., 2008) while the D,D-transpeptidases use the disaccharide-pentapeptide *N-*acetylglucosamine-*N-*acetylmuramic acid-L-Alanyl-D-Glutamyl-*meso*-diaminopimelyl-D-alanyl-D-alanine as their substrate (Tolufashe et al., 2020). Interactions between the peptide subunits of these nascent PG substrates with their relevant enzymes have been described (Cordillot et al., 2013; Erdemli et al., 2012; Fakhar et al., 2017; Lavollay et al., 2008; Mainardi et al., 2005), and it is generally assumed that the disaccharide component of the subunit interacts only with transglycosylases (Fibriansah et al., 2012; Mavrici et al., 2014) and thus are not relevant to the transpeptidases. However, evidence challenging this model is growing. Recent studies with the *Bacillus subtilis* L,D-transpeptidase LdtBs have suggested binding of the disaccharide component through a PG recognition domain, LysM, within LdtBs (Schanda et al., 2014). However, the mechanism of PG recognition may be different in *M.tb* since this LysM domain is absent in its L,D-transpeptidases.

In the current study, we investigate the interaction of PG substrate and Ldt_Mt2_. Using several interdisciplinary approaches, we have identified a new pocket in the Ldt_Mt2_ enzyme that binds PG saccharide moiety. We further elucidate the role of this pocket in recognition of ß-lactams in cooperativity with the catalytic site. These observations not only explain the mechanism for manifestation of physiological activity of Ldt_Mt2_, but also give insights into inhibition by ß-lactams that provide the context for structure-based design of anti-tubercular drugs.

## RESULTS

### A pocket remote from the catalytic site of Ldt_Mt2_ binds to peptidoglycan

The crystal structure of Ldt_Mt2_ was solved at 1.57Å resolution (**Table 1**). An electron density was observed in a pocket between the IgD2-YkuD domains, and a glucose molecule could be modeled into the electron density at 1.0 sigma (**Fig. 1A and S1**). This glucose molecule is likely to be part of a PG disaccharide moiety originating from the *E.coli* cell lysate during Ldt_Mt2_ purification. The sugar molecule is ensconced at the IgD2-YkuD domain interface in a pocket, which we referred to as the S-pocket, making several electrostatic interactions with residues R209, E168, R371, Y330 and A171 (**Fig. 1A**). Three residues M157, A171 and L391 stabilize the sugar through hydrophobic interactions. To provide additional evidence for the binding of PG substrates within the S-pocket, we performed ThermoFluor assays with N-Acetylmuramyl-L-alanyl-D-isoglutamine hydrate, a precursor of PG. A higher molar concentration of N-Acetylmuramyl-L-alanyl-D-isoglutamine gradually shifted the melting curve of Ldt_Mt2_ indicative of saturable binding behavior. A single R209E mutation in the S-pocket disrupted the binding of N-Acetylmuramyl-L-alanyl-D-isoglutamine with Ldt_Mt2_ (**Fig. 1B**).

**Table 1.**
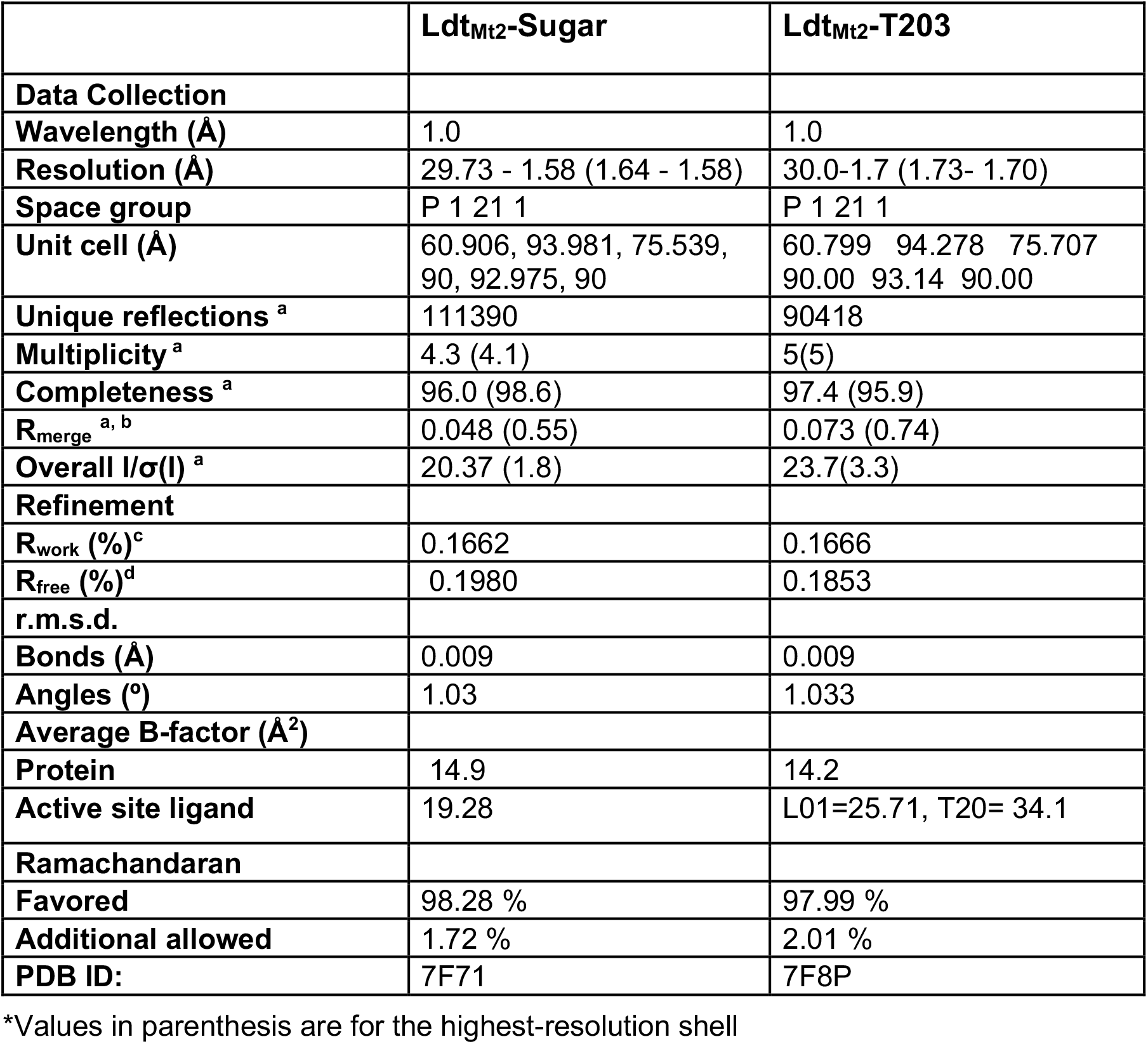
Data collection and refinement statistics

|  | **Ldt<sub>Mt2</sub>-Sugar** | **Ldt<sub>Mt2</sub>-T203** |
| --- | --- | --- |
| **Data Collection** |  |  |
| **Wavelength (Å)** | 1.0 | 1.0 |
| **Resolution (Å)** | 29.73 - 1.58 (1.64 - 1.58) | 30.0-1.7 (1.73- 1.70) |
| **Space group** | P 1 21 1 | P 1 21 1 |
| **Unit cell (Å)** | 60.906, 93.981, 75.539,<br>90, 92.975, 90 | 60.799 94.278 75.707<br>90.00 93.14 90.00 |
| **Unique reflections <sup>a</sup>** | 111390 | 90418 |
| **Multiplicity <sup>a</sup>** | 4.3 (4.1) | 5(5) |
| **Completeness <sup>a</sup>** | 96.0 (98.6) | 97.4 (95.9) |
| **R<sub>merge</sub> <sup>a, b</sup>** | 0.048 (0.55) | 0.073 (0.74) |
| **Overall I/σ(I) <sup>a</sup>** | 20.37 (1.8) | 23.7(3.3) |
| **Refinement** |  |  |
| **R<sub>work</sub> (%)<sup>c</sup>** | 0.1662 | 0.1666 |
| **R<sub>free</sub> (%)<sup>d</sup>** | 0.1980 | 0.1853 |
| **r.m.s.d.** |  |  |
| **Bonds (Å)** | 0.009 | 0.009 |
| **Angles (°)** | 1.03 | 1.033 |
| **Average B-factor (Å<sup>2</sup>)** |  |  |
| **Protein** | 14.9 | 14.2 |
| **Active site ligand** | 19.28 | L01=25.71, T20= 34.1 |
| **Ramachandaran** |  |  |
| **Favored** | 98.28 % | 97.99 % |
| **Additional allowed** | 1.72 % | 2.01 % |
| **PDB ID:** | 7F71 | 7F8P |

\*Values in parenthesis are for the highest-resolution shell

**Figure 1.**
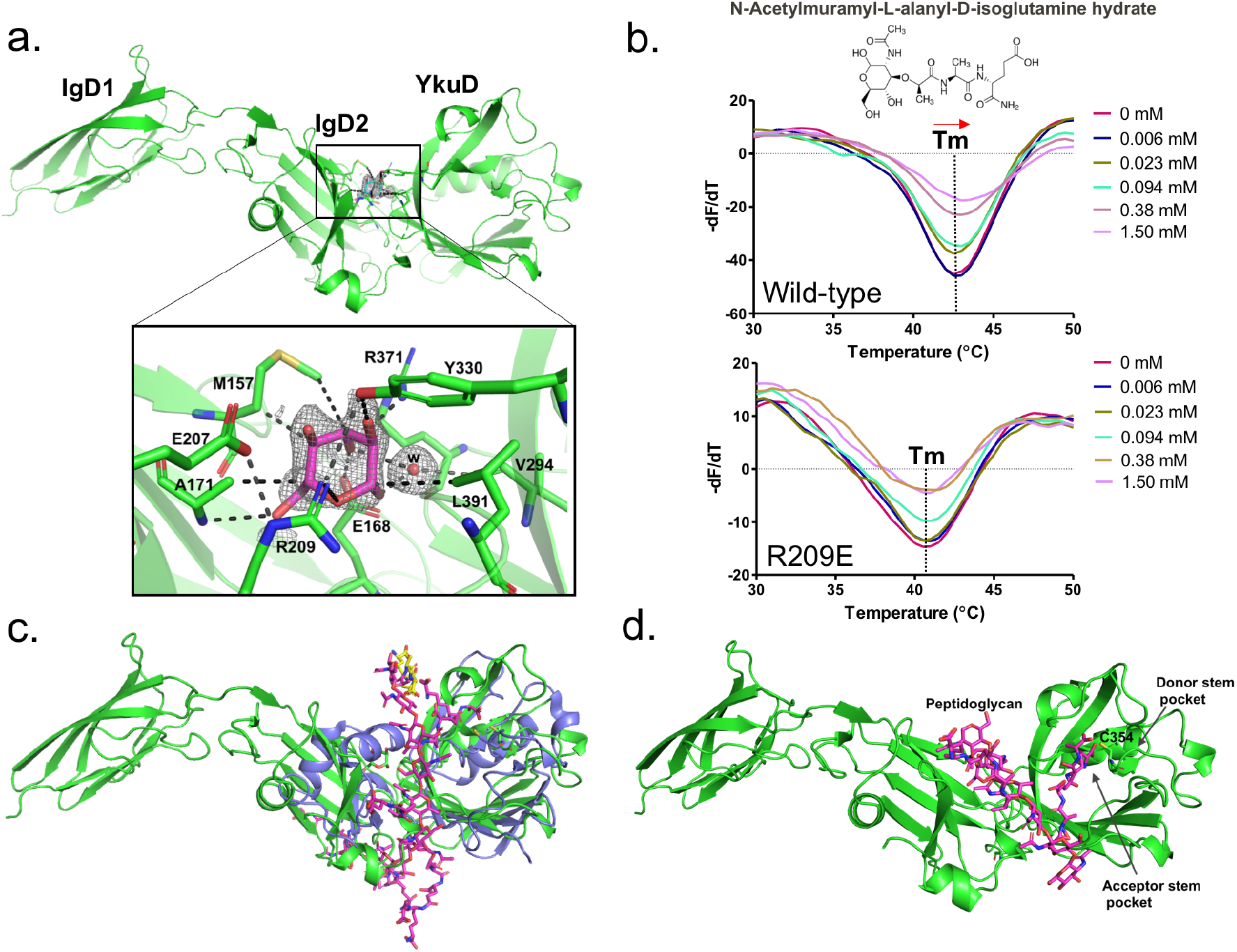
Binding studies of peptidoglycan with Ldt_Mt2_. (a) Crystal structure of Ldt_Mt2_ in complex with one glucose molecule. The inset shows the 2Fo-Fc omit map (contoured at 1.0σ) of glucose (cyan colour) modelled into the S-pocket of Ldt_Mt2_ in the crystal structure. (b) ThermoFluor assay for binding studies with the PG-precursor N-Acetylmuramyl-L-alanyl-D-isoglutamine hydrate with wild type Ldt_Mt2_ and the R209E mutant. The dotted line indicates the Tm, and a red arrow indicates the direction of thermal shift. Assays were performed in biological triplicates, and graphs were plotted by fitting the data on Sigmoidal dose-response (variable slope) equation in GraphPad Prism software. (c) Superposition of Ldt_Mt2_ (green) with PG-bound Ldt_BS_, the *Bacillus subtilis* L,D-transpeptidase (PDB ID: 2MTZ) (blue). (d) Modelling of peptidoglycan (pink color) into the Ldt_Mt2_ crystal structure (green).

An atomic model of the L,D-transpeptidase from *Bacillus subtilis* (LdtBS) in complex with nascent PG chain has been reported earlier (PDB ID: 2MTZ) (Schanda et al., 2014). Superposition of the structures of the Ldt_Mt2_-sugar complex with the Ldt_BS_-PG complex suggests that longer nascent PG chains thread across the S-pocket in between the IgD1-YkuD domains of Ldt_Mt2_ (**Fig. 1C**). Based on the structural details of PG binding in Ldt_BS_ and Ldt_Mt2_, a pentameric PG chain was computationally placed across the IgD1-YkuD domain interface encompassing the S-pocket. This computational modeling of a longer PG chain spatially aligns one of its tetrapeptide stem across an inner cavity of the catalytic site (**Fig. 1D**), similar to reports of carbapenem binding in the same position(Bianchet et al., 2017). This inner cavity of the catalytic site is proposed to bind the acceptor tetrapeptide stem, and the outer cavity to bind the donor tetrapeptide stem prior to their 3-3 transpeptide cross-linkage by Ldt_Mt2_ (Erdemli et al., 2012). Based on our crystal structure and modeling study, we propose that the S-pocket anchors the disaccharide moiety of one of the nascent PG chains prior to transpeptidation of tetrapeptide stems in the catalytic site.

### The S-pocket modulates ß-lactam hydrolysis activity

Our crystal structure reveals that the Ldt_Mt2_ enzyme is composed of three distinct domains as shown in figure 2A. As the S-pocket resides within the IgD2-YkuD domain interface, we investigated whether the S-pocket or different Ldt_Mt2_ domains play contributing role in the enzyme’s catalytic function. Due to the lack of tractable enzymatic assays with native PG substrates for observing physiological catalytic activity, we choose nitrocefin, a chromogenic ß-lactam, as a reporter substrate to assess the ß-lactam hydrolysis activity (Bianchet et al., 2017). To undertake this study, we expressed and purified fragments of Ldt_Mt2_ corresponding to IgD1, IgD2, IgD1-IgD2, IgD2-YkuD and YkuD domains. The full-length Ldt_Mt2_ holoenzyme showed a V_max_ of 0.23 μM/min and K_m_ of 16.32 μM in the nitrocefin hydrolysis assay (**Table 2**). Deletion of the IgD1 domain assessed by the IgD2-YkuD domain fragment resulted in no effect on the ß-lactam hydrolysis. However, deletion of IgD2 from the YkuD domain as assessed by the YkuD fragment alone led to a significant adverse effect on the nitrocefin hydrolysis activity with an increase in the Km value to 129 μM (an ~8-fold increase) and a decline in enzyme turn-over by ~10-fold (**Fig. 2B and Table 2**). This suggests an important role of S-pocket which is carried by the IgD2 domain in governing the catalytic activity of Ldt_Mt2_ enzyme. We further evaluated the role of the S-pocket by generating site-directed mutations at residues R209 and Y330 as they are situated within the S-pocket (**Fig. 2C**). The R209E mutant hydrolyzed nitrocefin with a K_m_ 428 μM that is ~26 fold higher than wild-type, while its V_max_ remained the same as wild-type. A high K_m_ value is an indicator of weak binding of the substrate with the enzyme, and shows that the enzyme would need a greater number of substrate molecules to achieve a V_max_. How a single R209E mutation in S-pocket would affect the ß-lactam hydrolysis activity in a catalytic site that is ~21 Å away remains a question; however, as shown below it is likely that the S-pocket has a role in modulating the catalytic activity of Ldt_Mt2_ enzyme allosterically.

**Table 2.**
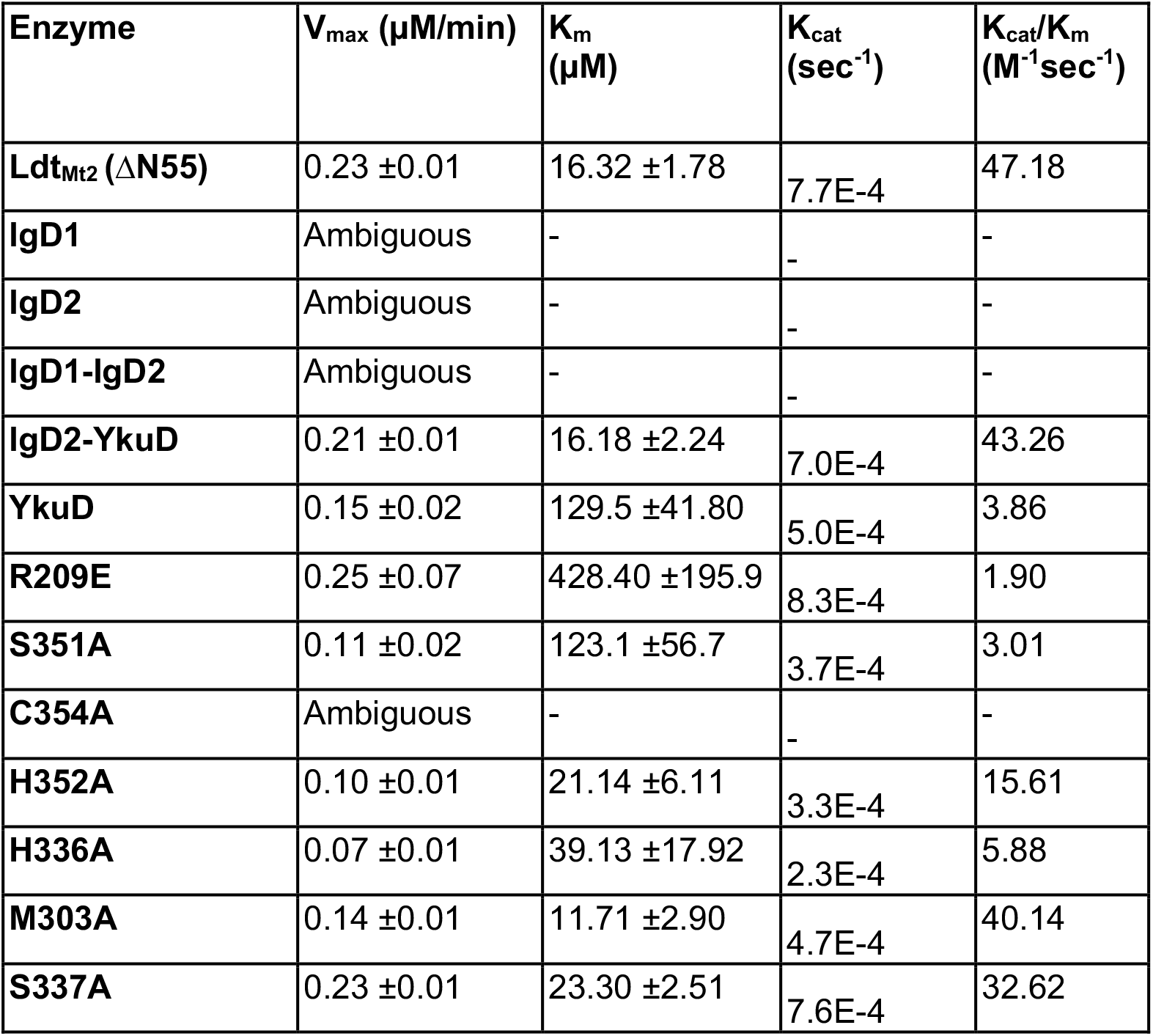
Kinetic parameters of ß-lactam hydrolysis by Ldt_Mt2_ and mutant proteins

**Figure 2.**
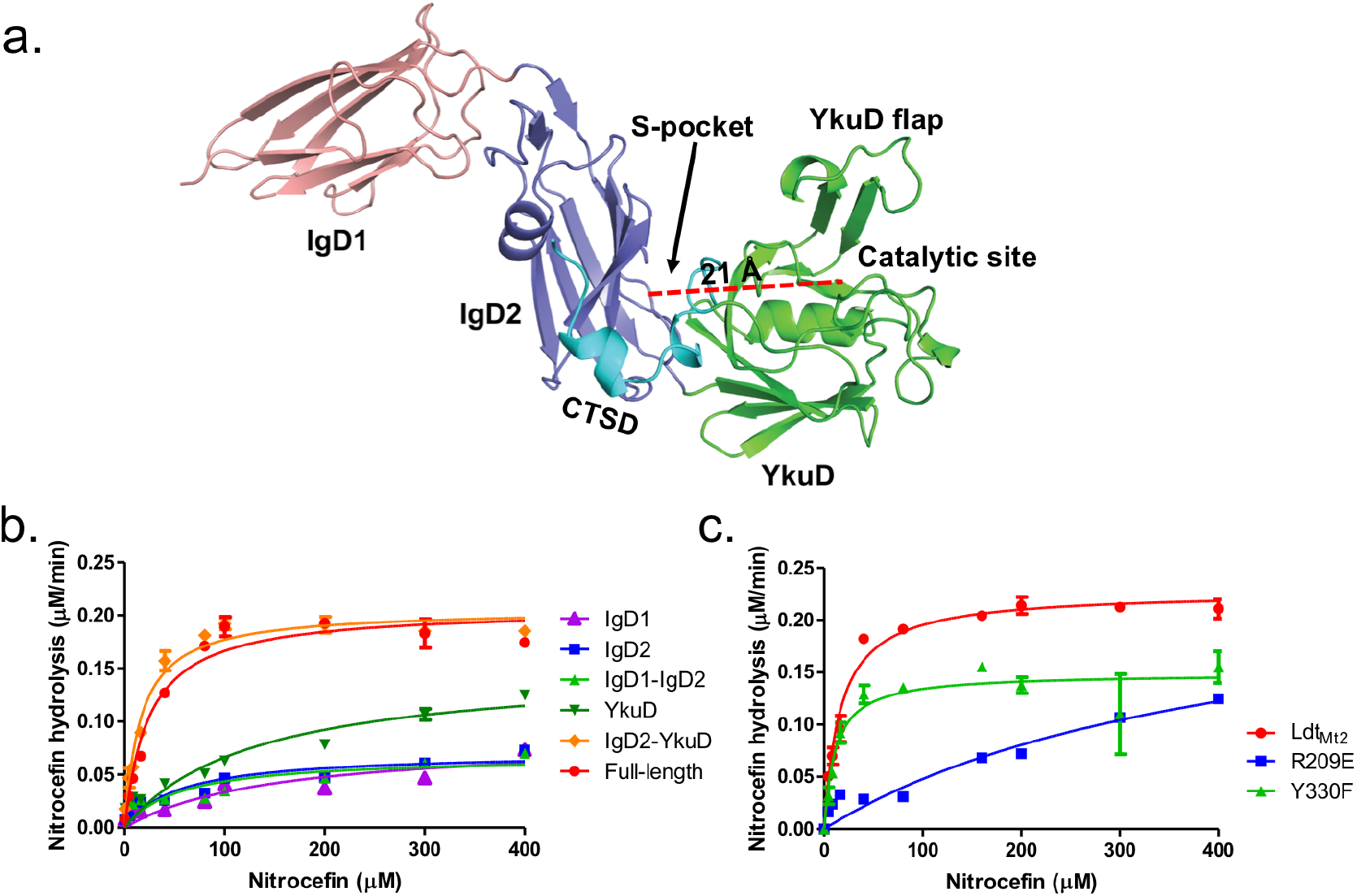
Role of the S-pocket in ß-lactam hydrolysis. (a) The structure of Ldt_Mt2_ with each domain highlighted: IgD1 (orange), IgD2 (blue), YkuD domain (green) and CTSD domain (cyan). A red dotted line demarcates the 21 Å distance between the S-pocket and the catalytic site. (b) Chromogenic nitrocefin hydrolysis activity of truncated Ldt_Mt2_ fragments corresponding to the IgD1, IgD2, IgD1-IgD2, YkuD, and IgD2-YkuD domains. (c) Chromogenic nitrocefin hydrolysis activity of wild-type Ldt_Mt2_, R209E, and Y330F mutants. Nitrocefin hydrolysis assays were performed in experimental duplicates and graphs were plotted in GraphPad Prism software by fitting the data on nonlinear regression curves using the Michaelis-Menten equation.

### The S-pocket cross-talks with the catalytic site to modulate ß-lactam hydrolysis

To evaluate the effects of mutations in the S-pocket on catalytic site activity ~21 Å away, we ran molecular dynamic (MD) simulations of the Ldt_Mt2_ wild-type and R209E mutant proteins. After 100 ns of MD simulations, structural and conformational changes were observed in the catalytic centre of YkuD domain including the catalytic triad residues C354, H336 and S337 and other residues in the catalytic site, namely, S351, M303 and W340. After the MD simulations in wild-type Ldt_Mt2_, H336 formed a hydrogen-bond interaction with the carbonyl oxygen of C354 as well as with the side chain hydroxyl group of S351. Unexpectedly, no hydrogen bond interaction was observed between H336-Nε1 and carbonyl oxygen of S337 (**Fig. 3A**) as this interaction was reported to be important for stabilizing the tautomer of H336 protonated at Nε1 (Erdemli et al., 2012).

**Figure 3.**
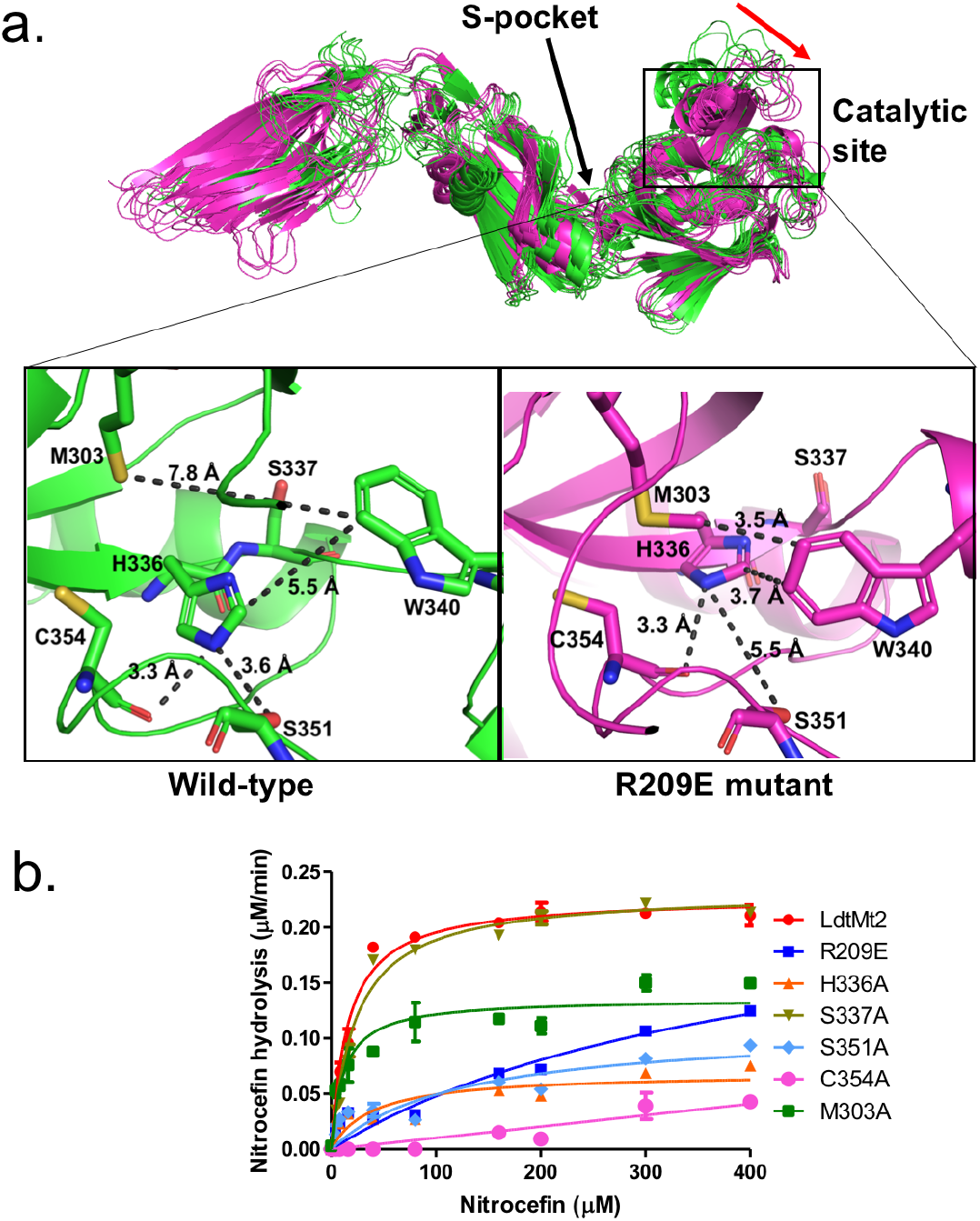
S-pocket crosstalk with the catalytic site of Ldt_Mt2_. (a) Molecular dynamic (MD) simulation of wild-type Ldt_Mt2_ (green) and the R209E mutant (pink). A red arrow indicates a shift in YkuD flap in R209E mutant during 100 ns of MD. The inset shows a detailed view of the catalytic site of the wild-type protein and R209E mutant after 75 ns of MD simulations. (b) Chromogenic nitrocefin hydrolysis activity of wild-type Ldt_Mt2_ and different mutants with alterations in both the S-pocket and catalytic site. Nitrocefin hydrolysis assays were performed in experimental duplicates and graphs were plotted in GraphPad Prism by fitting the data on nonlinear regression curves with the Michaelis-Menten equation.

When we assessed the MD simulated structure of the R209E mutant, no hydrogen bond interaction were found between H336 and the hydroxyl group of the S351 residue; however, the hydrogen bond interaction with carbonyl oxygen of C354 remained conserved. Additionally, the hydrogen bond interaction between the H336-Nε1 residue and the carbonyl oxygen of S337 residue was restored. Another W340 residue that resides outside the catalytic pocket comes closer to M303 and H336 through hydrophobic interactions to block the outer pocket of catalytic core. Such blockage of outer catalytic pocket would also hinder the dynamics of the YkuD flap, which has been reported to be important in ß-lactam binding(Fakhar et al., 2017). We also observed an overall difference in the flexibility state of YkuD flap region (300-330 residues) between the wild-type and R209E mutant structures after 100ns of MD simulation (**Fig. S2**).

To identify the functional relevance of conformational changes in the catalytic core residues upon MD simulations in both wild-type and R209E mutant structures, we performed *in-vitro* ß-lactam hydrolysis activity with site-directed mutants of the catalytic triad residues C354, H336 and S337 and other residues namely M303 and S351 that showed significant conformation changes upon R209E mutation in MD experiments. To our surprise, mutation of catalytic triad residue S337 to S337A did not disrupt the ß-lactam hydrolysis activity. The S337 residue was earlier reported as an important part of catalytic triad via stabilization of the protonated H336 tautomer (Erdemli et al., 2012). Mutation of C354 to C354A and H336 to H336A disrupted ß-lactam hydrolysis as expected. Additionally, mutation of the S351 residue to S351A disrupted ß-lactam hydrolysis activity, almost to the same degree as seen in the H336A mutant, with a ~15-fold decrease in enzyme turn-over (**Fig. 3B, Table 2**). In MD simulation runs with the wild-type structure, the S351 sidechain hydroxyl group forms hydrogen bond interaction with H336-Nε2, and this hydrogen bond interaction is absent in the R209E mutant structure (**Fig. 3A**). H336 is important for deprotonating the C354 sulphur to allow nucleophilic attack on carbonyl group of ß-lactam ring (Erdemli et al., 2012). We suggest that instead of the S337 residue, it is S351 that may form the catalytic triad together with C354 and H336 to stabilize the protonated H336 tautomer during ß-lactam binding and hydrolysis.

### Both the S-pocket and catalytic site participate in ß-lactam recognition

Among ß-lactams, the penicillin and cephalosporin classes are readily hydrolyzed by Ldt_Mt2_, while the carbapenem class inhibits Ldt_Mt2_ by irreversible acylation of C354 residue in the active site. In the current study, we used the carbapenem molecule, biapenem, to evaluate acylation of Ldt_Mt2_. The rate of acylation by biapenem was measured by monitoring a decrease in biapenem absorbance at 292nm wavelength. A single R209E mutation in the S-pocket completely disrupted biapenem-mediated acylation of the Ldt_Mt2_ enzyme (**Fig. 4A**). Mutation of catalytic residues C354, H336 and S351 also abrogated acylation with biapenem. These findings suggest that both the S-pocket and the catalytic center play important roles in driving acylation of the C354 catalytic residue by biapenem.

**Figure 4.**
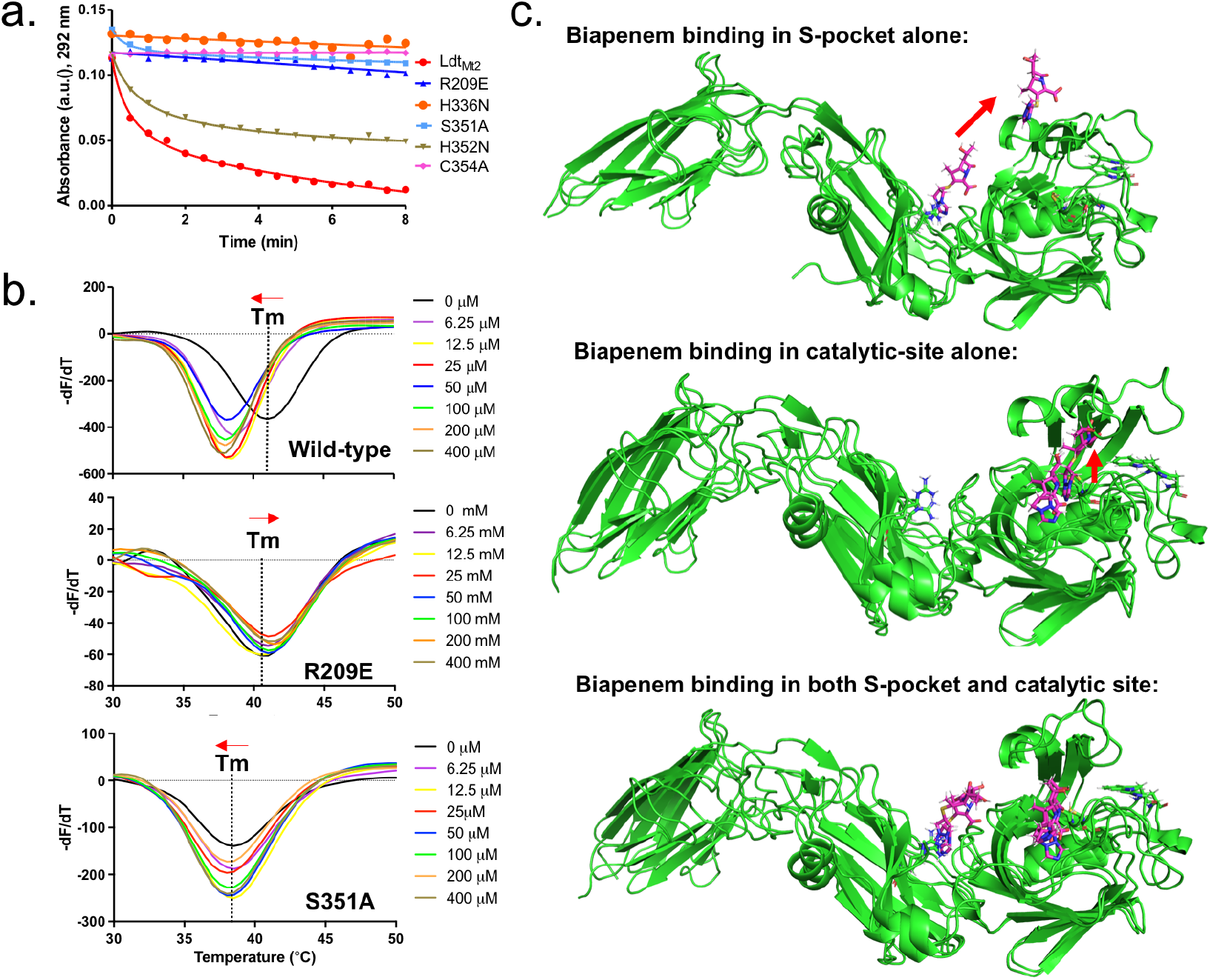
Role of the S-pocket and catalytic site in recognizing biapenem. (a) Acylation activity of biapenem with Ldt_Mt2_ and mutants was monitored at 292nm wavelength using UV–Visible spectrophotometry. Maximum absorbance spectra of biapenem was found at 292 nm that was used to monitor decrease in biapenem concentration upon acylation with the Ldt_Mt2_. The experiment was performed in biological triplicates to calculate the average values and standard deviations. Graphs were plotted in GraphPad Prism by fitting the data on a nonlinear regression curve with one-phase decay. (b) ThermoFluor assays for binding of biapenem with Ldt_Mt2_ and the R209E and S351 mutants. (c) Molecular dynamic simulations of Ldt_Mt2_ in complex with biapenem. Ldt_Mt2_ is represented in green and biapenem in pink. The red arrow indicates the movement of biapenem to a second position revealed by the MD simulations.

To further understand the role of the S-pocket and catalytic site in biapenem binding to Ldt_Mt2_, we performed ThermoFluor assays. Different amounts of biapenem (0-400μM) were titrated into 5.0 μM of Ldt_Mt2_ enzyme, and thermal shifts were measured at different drug concentrations. These studies revealed interesting observations: (1) increasing concentrations of biapenem led to a gradual change in melting temperature of Ldt_Mt2_ until it was fully saturated, and (2) biapenem binding decreased the melting temperature of protein (**Fig. 4B**). In the first observation, we found that Ldt_Mt2_-biapenem binding could be saturated only by enzyme:drug ratios as high as 1:80. This strongly suggests that biapenem saturates a surface of Ldt_Mt2_ through reversible, non-covalent interactions, as the covalent interactions have to be with a 1:1 molar ratio and are reversible. Beyond to its well-known covalent binding at the catalytic site (Kumar et al., 2017), these findings are consistent with non-covalent, saturable binding of biapenem to a second surface on Ldt_Mt2_. From the second observation, we conclude that biapenem binding destabilizes the protein possibly through structural changes. This structural destabilization may supersede the well-known structural changes in the YkuD flap at the catalytic site that are known to occur during ß-lactam binding and covalent reaction with the S^ϒ^ atom of C354 (Bianchet et al., 2017; Fakhar et al., 2017; Kim et al., 2013).

As the S-pocket mutant R209E exhibited diminished ß-lactam hydrolysis (**Fig. 3B**) and acylation by biapenem (**Fig. 4A**), we further analyzed the consequence of the R209E mutation on the physical binding of biapenem. In constrast to a the gradual decrease the thermal stability displayed by wild-type Ldt_Mt2_ upon biapenem binding, the R209E mutant showed only a subtle increase in Tm, and the saturating property of biapenem was virtually absent even at the highest concentration of 400μM (**Fig. 4B**). We conclude that the R209E mutation in the S-pocket hindered both non-covalent (as seen in **Fig. 4B**) as well as covalent interactions with biapenem (as seen in **Fig. 4A**). Additionally, as biapenem binds neglibly to the R209E mutant in contrast to wild-type Ldt_Mt2_, we did not observe decreases in the melting temperature of the R209E mutant with added biapenem as would be anticipated via catalytic site structural changes in the YkuD flap(Fakhar et al., 2017). This is further illustrated by our MD simulation results wherein the R209E mutation brings W340 residue closer to the YkuD flap residue M303 and active-site core residue H336 to block access to the outer pocket of the catalytic site (**Fig. 3A**). These R209E mutation-driven structural changes in the catalytic site and YkuD flap likely account for the inability of biapenem to bind to the R209E mutant of Ldt_Mt2_.

As structural changes occur in the catalytic site due to mutations in the S-pocket (**Fig. 3**), we hypothesized that catalytic site might also demonstrate an interplay with the S-pocket to indirectly influence non-covalent binding of biapenem. Binding studies were performed between biapenem and a catalytic mutant S351A using ThermoFluor assays. Indeed, the S351A mutant showed a significant decrease in its thermal stability (**Fig. 4B**). Moreover, varying concentrations of biapenem (0-400μM) with the S351A mutant did not induce any further significant thermal shift, indicative of neglible or insignificant physical binding of biapenem. Acylation with biapenem at the catalytic site was also diminished upon S351A mutation in Ldt_Mt2_ (**Fig. 4A**). These experimental observations indicate that mutations in catalytic site such as S351A disrupt both non-covalent as well as covalent binding of biapenem to Ldt_Mt2_.

### Two ß-lactams are recognized through cooperativity between the S-pocket and the catalytic site

From the experimental results with the wild-type, R209E and S351A mutants (**Fig. 4**), we conclude that biapenem has two modes of binding: (1) covalent binding and (2) saturable, non-covalent binding. In both of these binding modes, our data supports a dual role of both S-pocket and the catalytic site. As covalent binding is well known at the catalytic site, saturable reversible binding seems to be at a second surface of Ldt_Mt2_. From the experimental results it is quite evident that biapenem binding at both the sites can be controlled by the S-pocket and the catalytic site. Thus, we hypothesized that both the S-pocket and catalytic site may cooperate with each other in recognition of biapenem. Towards discovering the structural basis of cooperativity, we performed docking and molecular dynamic simulation studies of biapenem with Ldt_Mt2_ (**Fig. 4C**). A Ldt_Mt2_ crystal structure (PDB ID: 5DU7) that has the catalytic site within the YkuD flap in closed conformation was chosen for docking with biapenem as this structure will not allow the drug binding in the catalytic pocket. A grid was assigned for docking within a 60 Å radius of the catalytic site residue C354. We found that biapenem docked well within the S-pocket through its pyrazolo[1,2-a][1,2,4]triazolium R3 group with a binding energy of −6.3 kcal/mol.

Next, MD simulation experiments were further performed with Ldt_Mt2_ structures having biapenem docked (1) alone in S-pocket, (2) alone in catalytic pocket), (3) both in S-pocket and catalytic site. In the MD simulations with biapenem docked in S-pocket alone, the drug remained in the pocket for 9 ns of MD trajectory before exiting the pocket (**Fig. 4C**). Snapshots of different trajectories of biapenem in the S-pocket are shown in **Fig. S3.** In MD simulations with biapenem docked in the catalytic pocket alone, the ß-lactam core ring fluctuated at a distance of 3.7-7 Å from the S^ϒ^ atom of C354 during 5-40 ns (**Fig. 4C and S4**).

However, when biapenem molecules were docked in both the S-pocket and catalytic site simultaneously, the pyrazolo[1,2-a][1,2,4]triazolium R3 group of biapenem remained ensconced in S-pocket for 0-6 ns, made hydrophobic interactions with Y330 and L391 at 7-15 ns, and its pyrrolidine ring made additional *πl*-*πl* interactions with F330 at 18-28 ns while remaining in the S-pocket, before finally moving out towards the YkuD flap of the catalytic site (**Fig. 4C and S4A**). In the catalytic site over the simulation interval, biapenem movement fluctuated less this time, and its ß-lactam carbonyl oxygen atom remained oriented towards S^Y^ atom of C354 during 0-35 ns (see snapshots of biapenem trajectory in **Fig. S4B)**. Later, from 40 ns onwards in the MD trajectory, when biapenem at S-pocket came closer to catalytic site (as shown in **Fig. S4A**), the average distance between carbonyl oxygen and S^Y^ atom of C354 became 3.5 Å and Y308 made a hydrogen bond with G332. Thus, MD simulations suggest that biapenem binding across the S-pocket surface imposes stability in fluctuations of ß-lactam movement in the catalytic site and the ß-lactam ring carbonyl group maintains a close distance with the S^Y^ atom of C354 that favor a nucleophilic attack (see **Fig. S4** and **S5B**). These MD simulations together with our experimental data support a model in which two biapenem molecules are recognized cooperatively by both the S-pocket and the catalytic site, with bind -covalent, saturable binding and covalent binding to acylate C354, respectively.

### Binding patterns of various classes of ß-lactams in the S-pocket

As Ldt_Mt2_ binds various ß-lactams with variable affinities (Bianchet et al., 2017), we performed docking studies of various classes of ß-lactams with the S-pocket. Ampicillin and oxacillin from the penicillin class, cefotaxime from the cephalosporin class, and a new experimental drug, T203, from carbapenem class were chosen. The different ß-lactams showed binding with the S-pocket of Ldt_Mt2_ with variable energy scores using Autodock vina (**Table 3**). Ampicillin docked into the S-pocket with a binding energy of −7.1 kcal/mol with its R1-group tail 2-amino-2-phenylacetyl ensconced in the S-pocket through several electrostatic and hydrophobic interactions with the M157, E207, R209, R371 and Y330 residues (**Fig. 5A**). Another penicillin class member, oxacillin (a penicillinase-resistant penicillin), displayed the highest binding energy of −8.3 kcal/mol through its R1 group 5-methyl-3-phenyl-1,2-oxazole-4-carbonyl binding in the S-pocket (**Fig. S6**). Cefotaxime docked to the S-pocket with a binding score of −7.8 kcal/mol through R1-group tail thiozol-4yl (**Fig. 5B**). The new carbapenem T203 docked to the S-pocket with the least −6.9 kcal/mol binding with its R3 group 2-isopropoxy-2-oxoethyl (**Fig. 5C**), similar to the biapenem R3 group (**Fig. 4C**). The ß-lactam ring moieties of all of these ß-lactams were found to be free of any interactions with the S-pocket or surrounding residues, similar to biapenem. However, after 18-28 ns of MD simulation trajectory, the pyrrolidine ring of biapenem could make *πl*-*πl* interaction with F330 (**Fig. 4C and Fig. S4A**), and it is possible that similar late binding interactions may occur similarly with other the ß-lactams.

**Table 3.**
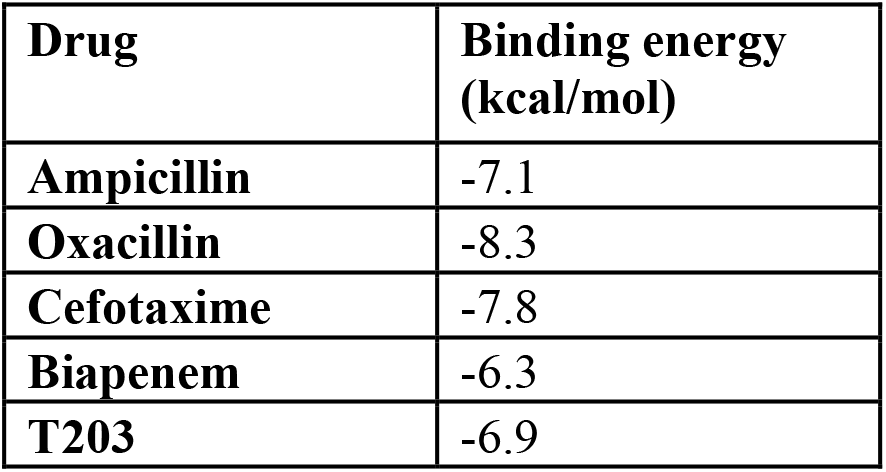
Binding energy of Ldt_Mt2_ with β-lactam compounds in kcal·mol^−1^ calculated by Autodock vina.

**Figure 5.**
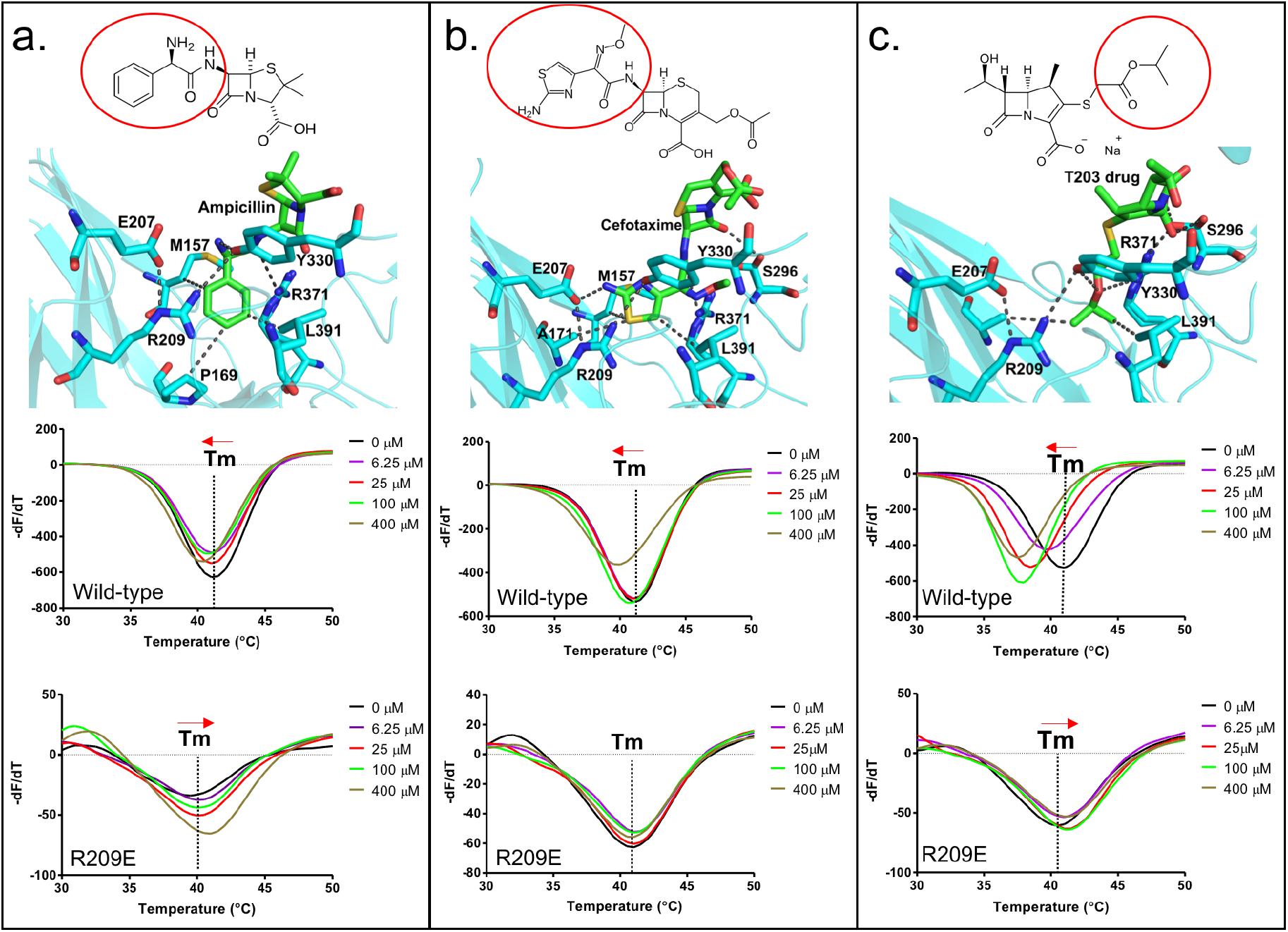
Binding of various classes of ß-lactams to the S-pocket. (a) Top: ampicillin (stick model in green) bound to the S-pocket (cyan) of Ldt_Mt2_ through its R1 group side-chain, 2-amino-2-phenylacetyl (red oval). Bottom: ThermoFluor assays for binding studies of ampicillin with wild-type Ldt_Mt2_ and the R209E mutant. (b) Top: cefotaxime (stick model in green) bound to the S-pocket (cyan) of Ldt_Mt2_ through its R1 group side-chain, thiozol-4yl (red oval). Bottom: ThermoFluor assays for binding studies of cefotaxime with wild-type Ldt_Mt2_ and the R209E mutant. (c) Top: the experimental carbapenem drug T203 (stick model in green) bound to the S-pocket (cyan) of Ldt_Mt2_ through its R3 group side-chain, 2-isopropoxy-2-oxoethyl (red circle). Bottom: ThermoFluor assays for binding studies of T203 drug with wild-type Ldt_Mt2_ and the R209E mutant.

ThermoFluor assays were also performed to investigate the binding behaviors of these additional ß-lactam class members with Ldt_Mt2_. Ampicillin, which has been reported to be readily hydrolyzed by Ldt_Mt2_ (Bianchet et al., 2017), showed a saturable binding behavior (**Fig. 5A**), but to a significantly lower degree than biapenem (**Fig. 4B**). Surprisingly, with ampicillin the R209E mutation in the S-pocket completely reversed the gradual thermal shift in Ldt_Mt2_ towards a higher Tm indicative of an increase in structural stability in the setting of clearly saturable binding (**Fig. 5A**). We interpret this to be consistent with reversible acylation of the C354 residue by ampicillin in addition to S-pocket binding. In support of this, a reversible acylation of the L,D-transpeptidase (Ldt_fm_ from *E.coli*) by ß-lactams in the catalytic site has been reported recently (Edoo et al., 2017; Zandi and Townsend, 2021). Oxacillin also showed a saturable binding with Ldt_Mt2_ (**Fig. S6**). With cefotaxime, the R209E mutation in the S-pocket strongly diminished saturable binding. And lastly, binding of new carbapenem drug T203 displayed a large thermal shift with Ldt_Mt2_ (**Fig. 5C**), similar to biapenem (**Fig. 4C**). We conclude that many ß-lactams (despite being weak or strong inhibitors of LtdMt2 activity) bind through the S-pocket with a saturable binding behavior; however, the carbapenem class brings maximum thermal destabilization in protein structure due to non-hydrolyzable covalent binding in catalytic site. Other classes of ß-lactam drugs, specifically the penicillins and cephalosporins, are known to be readily hydrolysed by Ldt_Mt2_ (Cordillot et al., 2013; Kumar et al., 2017).

### Structural basis of allosteric changes between the S-pocket and catalytic site

To further understand the high-resolution details of structural changes may that occur in Ldt_Mt2_ upon ß-lactam binding, the crystal structure of Ldt_Mt2_ was solved in complex with the new carbapenem drug T203 at a 1.7 Å resolution. Electron densities were observed in both the S-pocket and the outer cavity of catalytic pocket in the Ldt_Mt2_. Consistent with our docking results of T203 drug with Ldt_Mt2_ (**Fig. 5C**), the 2-oxoethyl side-chain of R3 group from T203 could be modelled into the electron density of the S-pocket. A second T203 drug was also modelled into the electron density map of the catalytic pocket. **Fig. 6A and 6B** show the 2Fo-Fc electron density map (contoured at 1.0σ) of T203 modelled in the S-pocket and catalytic site of the Ldt_Mt2_ in the crystal structure.

**Figure 6.**
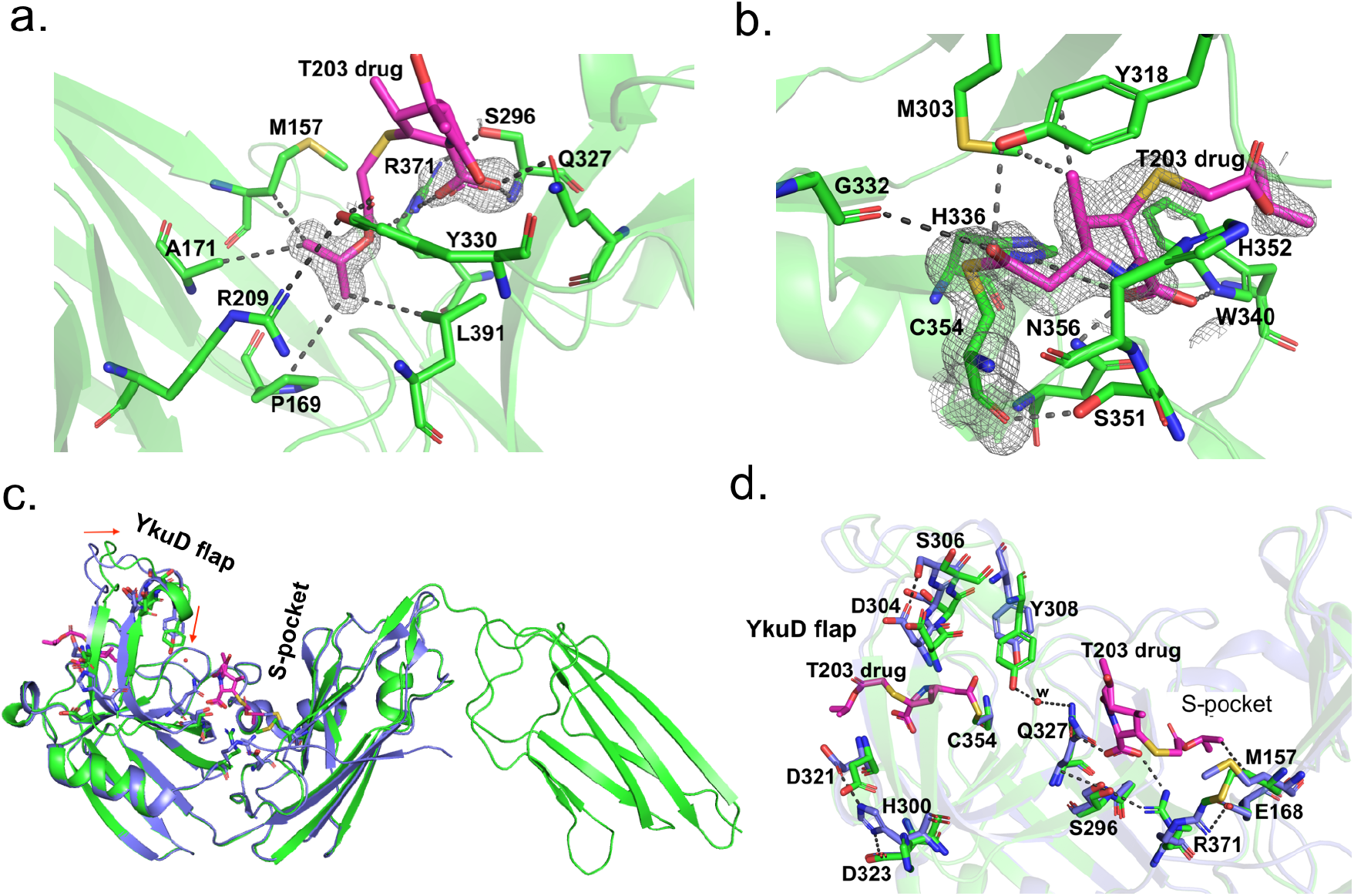
Structural studies of Ldt_Mt2_ with the experimental T203 carbapenem drug and allosteric conformation analyses. (a) The 2Fo-Fc map (contoured at 1.0σ) of the T203-R3 group side chain, 2-isopropoxy-2-oxoethyl (pink), modelled in the S-pocket of Ldt_Mt2_ in the crystal structure. (b) The 2Fo-Fc omit map (contoured at 1.0σ) of the full T203 structure (pink) modelled in the catalytic-site of Ldt_Mt2_ where it acylates the C354 residue of Ldt_Mt2_. (c) Superposition of the Ldt_Mt2_-T203 complex (green) with C354A catalytic mutant structure (PDB ID: 3TX4, blue). The red arrows indicate movements in YkuD flap upon T203 drug binding. (d) Residues that have undergone allosteric alterations upon T203 drug binding are shown with stick models. Ldt_Mt2_-T203 complex residues are represented in green and the C354A catalytic mutant in blue.

In the S-pocket of Ldt_Mt2_, the 2-oxoethyl sidechain of T203 drug is stabilized through hydrophobic interactions with the A171, M157, P169 and L390 residues (**Fig. 6A and S7A**). R371 makes an electrostatic interaction with the oxygen of the 2-oxoethyl moiety. No electron density was observed for the pyrrolidine ring of T203, while its carboxylic group fitted into an electron density making electrostatic interactions with backbone nitrogen of S296 and the guanidium side chain of R371. The modelling results of T203 into the electron density of the S-pocket were similar to the docking results of T203 drug and biapenem that also have their R3 group ensconced into the S-pocket with their pyrrolidine ring remaining free of any interaction with Ldt_Mt2_ (**Fig. 5**).

In the catalytic site, the T203 carbapenem interacts with the outer cavity at a covalent distance from the S^Y^ atom of C354 (**Fig. 6B and S7B**). The carbonyl oxygen of T203 makes hydrogen bond interactions with the hydroxyl group of Y318. The electron density for the R1-hydroxy ethyl group was not found, similar to three other related new carbapenems T206, T208 and T210 (Bianchet et al., 2017; Kumar et al., 2017). The methyl group of the pyrrolidine ring makes hydrophobic interaction with the phenyl ring of Y318. The amino N4 of the pyrrolidine ring makes electrostatic interactions with Nε2 of H336 and the backbone amide nitrogen of H352. The carboxyl group at C3 of the pyrrolidine ring makes hydrogen bond interactions with the side chains of W340 and N356. W340 also forms hydrophobic interactions with the 2-oxoethyl tail of T203.

We compared the structure of the Ldt_Mt2_-T203 complex with the C354A catalytic mutant structure (PDB ID: 3TX4) to seek alterations in conformation states of the enzyme around its catalytic site, YkuD flap, and S-pocket upon ß-lactam binding. We chose the catalytic mutant structure of Ldt_Mt2_ for structural comparison studies only because the wild-type enzyme usually binds ligands and/or substrates from its recombinant bacterial source during the purification steps(Erdemli et al., 2012), including in the current study. We observed that binding of the T203 drug introduces unique allosteric alterations in the salt bridge and hydrogen bond interactions spanning the entire distance from the S-pocket to the YkuD flap of the catalytic site. Upon T203 drug binding, the YkuD flap bends slightly towards the S-pocket (**Figure 6C**). In the S-pocket, the M157 side chain moves closer to the drug by 1.5 Å to make a hydrophobic interaction with the 2-oxoethyl tail of T203 drug (**Fig. 6D**). The R371 residue that was making salt bridge with E168 moves towards S296 through a hydrogen bond interaction and makes an additional ionic interaction with the carboxyl group of the T203 drug. The Q327 side chain that was previously producing a steric conflict with the carboxyl group of T203 drug moves away by a distance of 1.8 Å to make a water-mediated salt bridge with the hydroxyl group of Y308 that also moves down towards the S-pocket by a distance of 2.1 Å. The T203 drug binding induces an additional alteration in the YkuD flap by breaking the hydrogen bond interactions of H300 with D323 as well as D321 and alse the interactions between D304 and S306. Breaking of these hydrogen bond interactions possibly relaxes the YkuD flap, enabling it to tilt towards the S-pocket mediated by new water-mediated salt bridge between Y308 and Q327. Alterations in the dynamics of the YkuD flap was also observed in MD simulations with the R209E mutant and by biapenem binding in S-pocket (**Fig. 3A and Fig. 4**).

In summary, the evidence supporting the identity of S-pocket as allosteric site includes its distance of 21 Å from the catalytic residue C354, the observation of saturable binding by ß-lactam drugs in addition to covalent binding, and lastly, the substantial conformational alterations between the S-pocket and catalytic site upon ß-lactam binding. Mutational changes in the S-pocket or the catalytic pocket nullifies all the allosteric communications that are otherwise important in cooperative binding of dual ß-lactams. The consequences of these mutational changes were confirmed by ß-lactam hydrolysis assays (**Fig. 2C & 3B**), acylation by biapenem (**Fig. 4A**) and ThermoFluor assays (**Fig. 4B & 5**) with different class of ß-lactams.

## DISCUSSION

In addition to the role of L,D-transpeptidases in remodelling the PG in non-replicating *M. tb* (Lavollay et al., 2008), this enzyme class is responsible for the resistance of *M.tb* to most ß-lactam drugs, except carbapenems (Cordillot et al., 2013; Gupta et al., 2010). The molecular mechanisms and physiological function of L,D-transpeptidases and the basis for their genetic susceptibility to selective ß-lactams remains incompletely understood. In this study we reveal important aspects of the physiological function of the *M.tb* L,D-transpeptidase enzyme, Ldt_Mt2_, identify a new PG disaccharide moiety binding pocket (named the S-pocket), and describe the S-pocket’s role in allosteric modulation of the transpeptidase active site. Additionally we observe that various ß-lactams bind to the S-pocket through their tail regions to bring about allosteric changes which predispose the catalytic site for covalent inactivation by a second ß-lactam. Based on our findings, we propose a mechanism of allosteric communication between the S-pocket and the catalytic site in facilitating dual ß-lactam and/or dual PG substrate binding in Ldt_Mt2_ (**Figure 7**).

**Figure 7.**
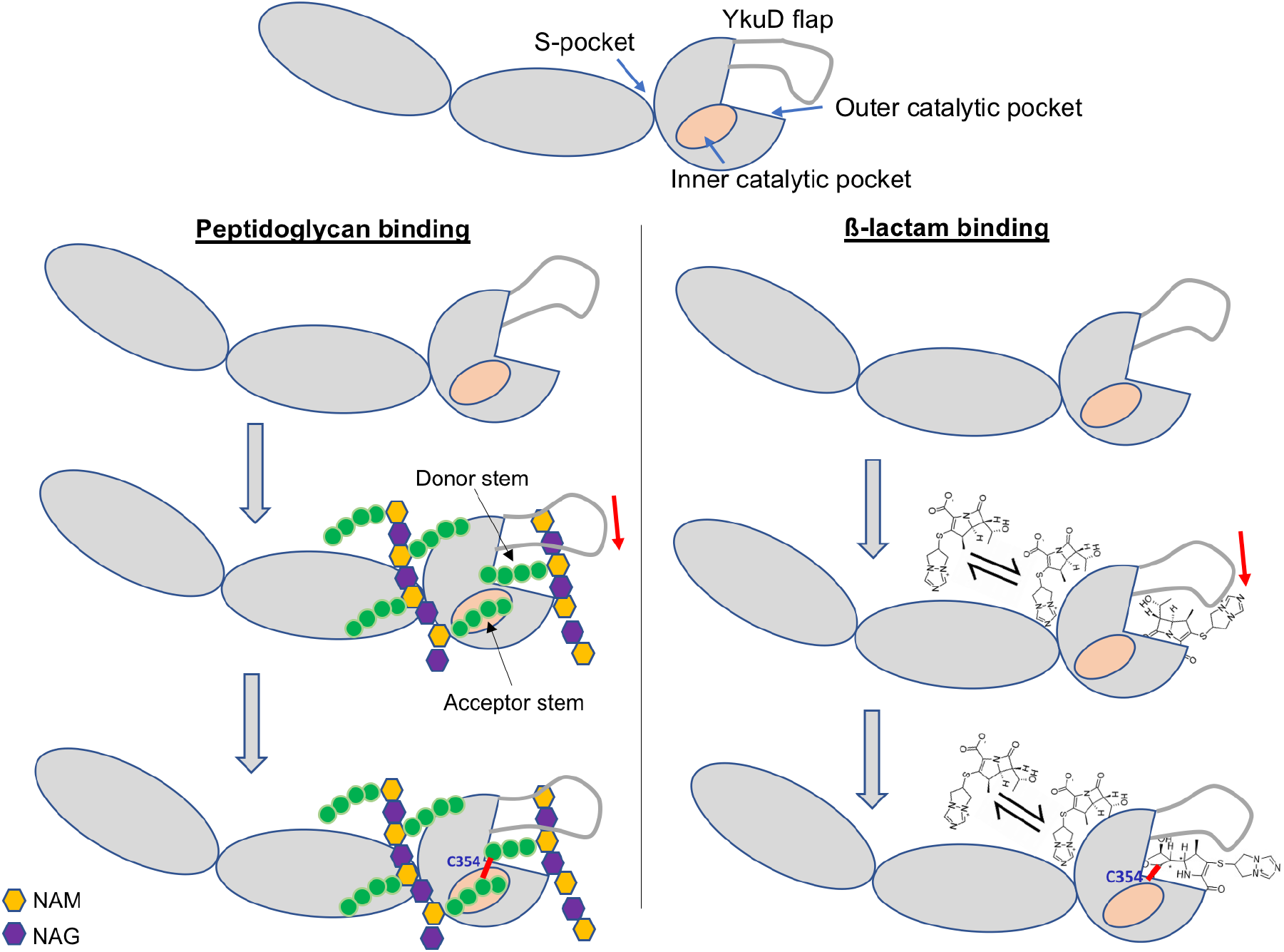
Cartoon model showing the mechanism of recognition of dual PG substrates and/or dual ß-lactam drugs across the S-pocket and catalytic site of Ldt_Mt2_. A small red line in the figure indicates a covalent bond between donor and acceptor stem peptides of PG or covalent bond between C354 and ß-lactam.

*M.tb* contains several paralogs of L,D-transpeptidases, namely LdtMt1, Ldt_Mt2_, Ldt_Mt3_, Ldt_Mt4_ and Ldt_Mt5_(Gupta et al., 2010). Crystal structures of Ldt_Mt1_(Correale et al., 2013), Ldt_Mt2_ (Erdemli et al., 2012), Ldt_Mt3_ (Libreros-Zuniga et al., 2019) and Ldt_Mt5_ (Brammer Basta et al., 2015) have been solved and reported to date. All of these paralogs contain a pocket similar to the S-pocket found in Ldt_Mt2_. Corresponding to the R209 residue position in Ldt_Mt2_ S-pocket, Ldt_Mt1_ has R25, Ldt_Mt3_ has Q66, and Ldt_Mt5_ has H219, and each of these putative S-pocket amino acids have similar basic charge properties (**Figure S8**). Moreover, superposition of the crystal structure of these paralogs with Ldt_Mt2_-sugar complex places the PG sugar moiety within the S-pocket. We suggest a common S-pocket-mediated allosteric mechanism in all of the L,D-transpeptidases in *M.tb*; however, the rate of transpeptidation may differ depending upon structural differences in their respective YkuD flaps, S-pockets and catalytic sites.

Bases on our crystal structure and modelling studies, we propose that prior to the 3-3 transpeptidation between the donor and acceptor PG stem peptides, the acceptor PG sugar moiety chain is anchored across the IgD1-YkuD domains interface to the S-pocket of Ldt_Mt2_. PG sugar chain anchoring has been observed in L,D-transpeptidase of *Bacillus subtilis* through a PG recognition domain LysM (Schanda et al., 2014). The LysM domain binds a sugar moiety of the PG precursor, and the tetrapeptide branch (acceptor stem) contacts the catalytic cysteine residue through the inner cavity of the catalytic domain. Another PG binding enzyme lysostaphin from *Staphylococcus simulans* has a PG anchoring domain, SH3b, while its catalytic domain cleaves PG stem cross-bridge (Mitkowski et al., 2019). Upon anchoring of the PG sugar moiety chain within the S-pocket in Ldt_Mt2_, its acceptor stem peptide binds to the inner pocket of the enzyme’s catalytic domain, and the donor stem binds to the outer cavity close to the C354 residue, interactions that foster formation of the 3-3 transpeptide linkage (Bianchet et al., 2017; Erdemli et al., 2012; Fakhar et al., 2017). We hypothesize that, prior to 3-3 transpeptide linkage, both S-pocket and catalytic site may work in cooperativity to facilitate synchronous binding of two PG substrates (donor and acceptor substrates); however, this requires experimental validation using nascent PG substrates that are beyond the scope of our study. Nevertheless, in support of our proposed model, we have tested our cooperativity hypothesis on ß-lactam binding and hydrolysis activity in Ldt_Mt2_ and found results that support the model.

Ldt_Mt2_ plays a major role in the resistance of *M.tb* to ß-lactam class of drugs (Cordillot et al., 2013; Dubee et al., 2012; Gupta et al., 2010; Lavollay et al., 2008; Mainardi et al., 2005). Among the ß-lactam class of drugs, penicillins and cephalosporins are readily hydrolyzed by this enzyme, while carbapenems are potent Ldt_Mt2_ inhibitors (Bianchet et al., 2017). Our findings reveal the role of both the S-pocket and the catalytic site in regulating ß-lactam hydrolysis and inhibition by the carbapenem class. We demonstrate an allosteric cooperativity between the S-pocket and the catalytic site in the dual recognition of carbapenem drugs, with the former one binding the carbapenem frug non-covalently with a saturable binding and the latter one covalently through irreversible acylation of C354. A similar ß-lactam binding mechanism has been observed in penicillin-binding protein 2a (PBP2a) from *Streptococcus aureus* where one molecule of ß-lactam occupies an allosteric site (with a saturable binding behavior) 60 Å away culminating into the allosteric conformational changes in PBP2a with the opening of the active site and covalent binding with a ß-lactam molecule(Otero et al., 2013). During the ß-lactam binding process, Ldt_Mt2_ occupies one molecule of ß-lactam at a reversible binding site (the S-pocket) 21 Å away from the catalytic site, and this interaction stimulates allosteric conformational changes across the YkuD catalytic flap to drive acylation by a second ß-lactam molecule in the catalytic pocket. We find the role of catalytic site equally important in stimulating reversible binding of ß-lactam in the S-pocket. Thus there we observe bi-directional cooperativity between the S-pocket and the catalytic site in binding dual ß-lactams, and the same mechanism may applied to dual PG substrate binding. The role of differential dynamics by the YkuD flap in ß-lactam- and substrate-binding by the catalytic site have been demonstrated earlier by MD simulations (Fakhar et al., 2017); however, the role of the YkuD flap in dual ß-lactam binding by the S-pocket and the catalytic site is demonstrated for the first time by this study. Several allosteric alterations mediated by new water-mediated salt-bridges or breakage of pre-existing ionic interactions contribute to the cumulative dynamics of the YkuD flap during dual ß-lactam binding in Ldt_Mt2_.

We find that various ß-lactams bind to the S-pocket of Ldt_Mt2_ through their tail regions, either through their R1 or R3 groups. As we found the docking scores of ampicillin, oxacillin, and cefotaxime to be higher than those of carbapenems, the interactions of these R1 or R3 groups with the S-pocket appears to play critical role in the initiation of S-pocket binding, irrespective of fate of ß-lactams in catalytic site. This discovery of a novel mechanism of ß-lactam binding in Ldt_Mt2_ reveals important new parameters in the development of novel ß-lactams for *M. tb*, and highlights the importance of the respective R1 and R3 side chains to both occupy the S-pocket and modulate strong inhibition at the catalytic site.

## SIGNIFICANCE

Biosynthesis of bacterial cell wall peptidoglycan (PG) is inhibited by the ß-lactam class of antibiotics. *Mycobacterium tuberculosis* susceptibility to ß-lactams is subclass specific as the carbapenems, but not the penicillins and cephalosporins, exhibit potent activity against this mycobacteria through effective inhibition of its L,D-transpeptidases, which catalyses 3-3 transpeptidation reaction in the biosynthesis of PG. A better understanding of L,D-transpeptidase function and mechanism of binding with natural substrate and various ß-lactams can provide insight necessary to leverage L,D-transpeptidases as targets for drug development. Based on our structural, biophysical and biochemical data, we identify a new PG disaccharide moiety binding pocket (named as S-pocket) at a distance of 21 Å from the catalytic site in the L,D-transpeptidase Ldt_Mt2_. This new site recognizes ß-lactams and modulate their hydrolysis. Our experimental and computational studies identify a allosteric cooperativity between S-pocket and the catalytic site in recognising dual ß-lactams, wherein ß-lactams bind S-pocket with saturable binding behaviour. Our crystallographic studies further reveal the high-resolution details of allosteric alterations that span across the S-pocket and catalytic site during dual ß-lactam binding. Identification of a cooperativity between S-pocket and catalytic site also represents a valuable case to investigate recognition of natural substrates prior to 3-3 transpeptide reaction. A model summarizing the molecular mechanism of two ß-lactams and/ substrate recognition is proposed based on our structural and biochemical data.

## MATERIALS & METHODS

### Cloning and site-directed mutagenesis

DNA sequences encoding Ldt_Mt2_-Δ42, Ldt_Mt2_-Δ55, IgD1 (50-145 aa residues), IgD2 (150-250 aa), IgD1-IgD2 (50-250 aa) and YkuD domain (250-408) and CTSD deletion mutant Ldt_Mt2_ 42-384 were cloned in pET28a vector to express the protein with *N*-terminal His_6_-tag that is cleavable by Tobacco Etch Virus (TEV) protease. Single amino acid substitutions of Ldt_Mt2_-Δ55 were constructed by site-directed mutagenesis for the following mutations: R209E, C354A, H352A, H336A, M303A, S337A and S351A as described(Bianchet et al., 2017). Primers used to clone different fragments of Ldt_Mt2_ using H37Rv chromosomal DNA are listed here:

IgD1 domain-Forward primer: attgccatatgaagggcacgccgttcgccgatc

IgD1 domain-Reverse primer: caatactcgagttaggtctggaaggtcagctggcg

IgD2 domain-Forward primer: attgccatatgacctgaccatgccctacgtcat

IgD2 domain-Reverse primer: caatactcgagttagccgatggtgaagtgcgtctg

IgD1+ IgD2 domain-Forward primer: attgccatatgaagggcacgccgttcgccgatc

IgD1+ IgD2 domain-Reverse primer: caatactcgagttagccgatggtgaagtgcgtctg

YkuD domain- Forward primer: attgccatatgggcgacgaggtgatcgcgacc

YkuD domain- Reverse primer: caatactcgagttacgccttggcgttaccggc

### Protein expression and purification

Mutants and different fragments of Ldt_Mt2_ were expressed and purified as reported earlier(Bianchet et al., 2017). In detail, Ldt_Mt2_-ΔN55 was transformed in chemical competent *E. coli* BL21δε3 (NEB labs). A single colony of transformed cells was inoculated in 50mL of Luria-Bertani (LB) media supplemented with ampicillin (100 μg/mL) before growing overnight (O/N) at 37°C in an incubator shaker. The O/N culture was used to inoculate secondary culture in LB media to grow at 37°C until the optical density at 600 nm reached ~0.6-0.8. At this stage, temperature was lowered to 16°C in the incubator shaker before inducing the protein expression with 0.5 mM of isopropyl-1-thio-β-galactoside (IPTG). The secondary culture grown O/N. The culture was harvested and the cell pellet was resuspended in lysis buffer (50 m*M* Tris buffer pH 7.5, 400 m*M* NaCl, 10% glycerol, 1.0 mM Dithiothreitol (DTT) and 1.0mM Phenylmethylsulfonyl fluoride (PMSF). 0.5mg/ml lysozyme was added into the resuspended cells to allow cell lysis at 4°C for 30 minutes. Resuspended cells were further lysed by ultrasonication at 4 °C with a pulse rate of 15 second ON/OFF. Whole cell lysate was centrifuged at 10,000g for 45 minutes and the supernatant was loaded onto Ni-NTA column (Qiagen, Germany). The unbound protein was washed with washing buffer (50 m*M* Tris buffer pH 7.5, 400 m*M* NaCl, 10% glycerol, 1.0mM DTT, 0.1 mM PMSF) and the protein was eluted with elution buffer (50 m*M* Tris buffer pH 8.0, 400 m*M* NaCl, 1.0mM DTT, 0.1 mM PMSF and 500 m*M* imidazole). The His6-tag of the protein was removed by TEV protease during overnight dialysis against the buffer 50 m*M* Tris pH 8.0, 150 m*M* NaCl, and 1.0mM DTT at 4°C. The dialyzed protein was passed through Ni-NTA column and the His6-tag-removed protein was collected in flow-through. Protein was further purified using superdex 10/300 column on ÄKTA™ pure 25. The purified protein was concentrated to 20 mg/ml as measured by nanodrop at 280nm wavelength. The purity of protein was checked by 12% SDS-PAGE. All other truncation and mutants of Ldt_Mt2_ were also purified by same protocol as above, however their His6-tag was not removed.

### ThermoFluor assays

The proteins Ldt_Mt2_-Δ55, R209E and S351A were with initial stocks of 11.5 μM, 14.0 μM and 21 μM respectively in the 50 mM Tris buffer pH 8.0, 150mM NaCl, 1 mM DTT. 5,000x of SYPRO™ Orange (Invitrogen) was diluted to 50x in water. 5 μM of proteins and 3x of SYPRO™ Orange were pipetted into a 96-well PCR plate (BioRad, MicroAmp Fast 96-Well Reaction plate, 0.1mL) with 50μl total volume in the well. Fluorescence data was collected on BioRad StepOnePlus Real-Time PCR System using the software StepOne software v2.3. ROX (SYPRO Orange) was selected as a reporter dye and none for passive reference in the software. The temperature was held for 1 min per degree from 25 to 65°C. Melting temperature (Tm) and differential fluorescence (-dF/dT) values were calculated by fitting the data on Sigmoidal dose-response (variable slope) equation in GraphPad Prism software. Experiments were performed in biological triplicates.

### Nitrocefin hydrolysis assays

Nitrocefin (Calbiochem) with a range of 1-400 μM was used as a substrate for quantifying the rate of ß-lactam hydrolysis by different Ldt_Mt2_ fragments and mutants. A 100μl reaction mixture containing 5 μM enzyme in 25 mM HEPES–MES–Tris-Phosphate buffer, 300 mM NaCl, pH 6.0, was incubated at 25°C. Nitrocefin hydrolysis was measured at 496 nm on BioRad microplate reader and the absorbance data were converted to μM/minute using Beer’s Law (*e* = 20,500 M^−1^ cm^−1^ for hydrolyzed nitrocefin; *L* = 0.5 cm). The rate constants, Vmax and Km were calculated by fitting the data on nonlinear regression curve with Michaelis-Menten equation.

### Biapenem acylation assays

The acylation of biapenem with Ldt_Mt2_ and mutants was determined by measuring the reduction in absorbance of biapenem at 292nm wavelength using UV–Visible spectrophotometry. A 100μl reaction mixture containing 50μM enzyme, 50μM Biapenem, 25mM tris buffer pH 7.5 was incubated at 15°C and endpoint absorbance was recorded at 30s intervals for 8 minutes. Rate constant (K) of biapenem acylation was calculated by fitting the data on a nonlinear regression curve with one-phase decay. Experiment was performed in biological triplicates to calculate standard deviation and average values.

### Protein Crystallization

Purified Ldt_Mt2_ (fragment ΔN55) was crystallized with the same conditions has reported earlier(Bianchet et al., 2017). Crystals were grown by hanging drop vapor diffusion method in 20% 5000MME and 200 mM ammonium sulphate condition. For Ldt_Mt2_-T203 complex, crystals were soaked with 2mM of T203 drug overnight before being cryo-protected in 20% 5000 MME, 30% glycerol and 120 mM ammonium sulphate before flash freezing in liquid nitrogen.

### Crystal diffraction, data collection and structure determination

The crystals were diffracted at 100K temperature at a wavelength of 1.0 Å on beamline 19-ID at the Advanced Photon Source (Argonne National Laboratory). The diffraction data were recorded on an ADSC Quantum 315r CCD detector and processed with the HKL3000 software.(Minor et al., 2006) The crystal structures of Ldt_Mt2_-sugar complex at a highest resolution of 1.58 Å and Ldt_Mt2_-T203 complex at 1.7 Å resolution were solved by molecular replacement method using *PHENIX* suite of program(Liebschner et al., 2019) using the coordinates of Ldt_Mt2_ (PDB ID: 5DU7) as a search model. The initial structures were subjected to crystallographic refinement with *phenix.refine(Afonine et al., 2012)* from the *PHENIX* suite of programs. Structures were rebuilt with COOT(Emsley and Cowtan, 2004) to fit the electron density map. Structure validation was done using Molprobity(Williams et al., 2018). The R values of refined structures (**Table 1**) are well within the range of typical resolution. Omit maps for ligands in the structures were created from map coefficient using *PHENIX* suite of programs. Figures were prepared using PyMOL Molecular Graphics System, Version 1.5.0.4 Schrödinger, LLC.

### Docking studies

Autodock vina(Trott and Olson, 2010) was used for docking studies. Ldt_Mt2_ (PDB ID: 5DU7) with closed active-site loop was used for docking studies with different ß-lactam ligands namely Ampicillin, Cefotaxime, Biapenem & T203. The grid for docking was assigned nearby the C354 residue so as to allow the ligands bind in close proximity (within 60 Å covering the area of PG- pocket).

### Molecular dynamics studies

Explicit water (TIP3P) MD simulations of Ldt_Mt2_, R209E mutant and Ldt_Mt2_-biapenem complex were carried out with AMBER16 employing ff03 force field(Duan et al., 2003). Leap module of AMBER16 was used for setting up initial structures. All solvated structures were energy minimized to prevent steric clashes. System were heated using Langevin dynamics from 10 to 300 K at NPT ensemble with a positional restraint of 5 kcal/mol/Å2. Positional restraint was released gradually in the next two steps i.e. 3 kcal/mol/Å2 in 1st step and then 1 kcal/mol/Å2 in 2nd step. Finally, the production runs (Ldt_Mt2_ = 100 ns, R209E = 100 ns, Ldt_Mt2__Bia-site_1 = 75 ns, Ldt_Mt2__Bia_site_2 = 75 ns, Ldt_Mt2__Bia_site_1_2 = 75 ns) were carried out at NPT ensemble by integrating the Newtonian equation of motion at every 2 fs. Trajectories were analysed using cpptraj module of AMBER16.

### Accession codes

Coordinates and structure factors of both Ldt_Mt2_-sugar complex and Ldt_Mt2_-T203 complex have been deposited in the PDB under the accession codes 7F71, 7F8P

## ACKNOWLEDGEMENTS

T203 drug was a kind gift from Dr. Joel S. Freundlich at Rutgers University Medical School. We thank personnel at Argonne National Laboratory for data collection of protein crystals. This study was supported by the funding from Department of Biotechnology, Government of India (BT-RLF/Re-entry/68/2017), SERB-Core Research Grant (CRG/2019/005079), Indian Council for Medical Research (ICMR) Adhoc grant (BMS/ADHOC/10/2019-20) and UGC-start-up grant (F.30-520/2020-BSR) to P.K. This study was also supported by NIH awards R33 AI111739 and R21 AI137720 to GL. This research was supported in part by PL-Grid Infrastructure. Computations were performed at Academic Computer Centre Cyfronet AGH.

## AUTHOR CONTRIBUTIONS

PK (study conceptualization and design, protein-drug interactions, protein crystallization, data collection and structural studies, data analysis, manuscript preparation), GL (study conceptualization and design, cloning and site-directed mutagenesis, manuscript preparation), WRB (data analysis and manuscript preparation), NA (Cloning & site-directed mutagenesis, protein expression & purification, ThermoFluor assays, data analysis), SK (computational studies), VC (protein expression and purification, nitrocefin hydrolysis and acylation assays with biapenem), KS (computational studies and data analysis), PJ (data analysis), CKB (data analysis and manuscript preparation). All authors contributed to the final draft of the manuscript. The authors declare no competing financial interests.

---

**Figure S1:**
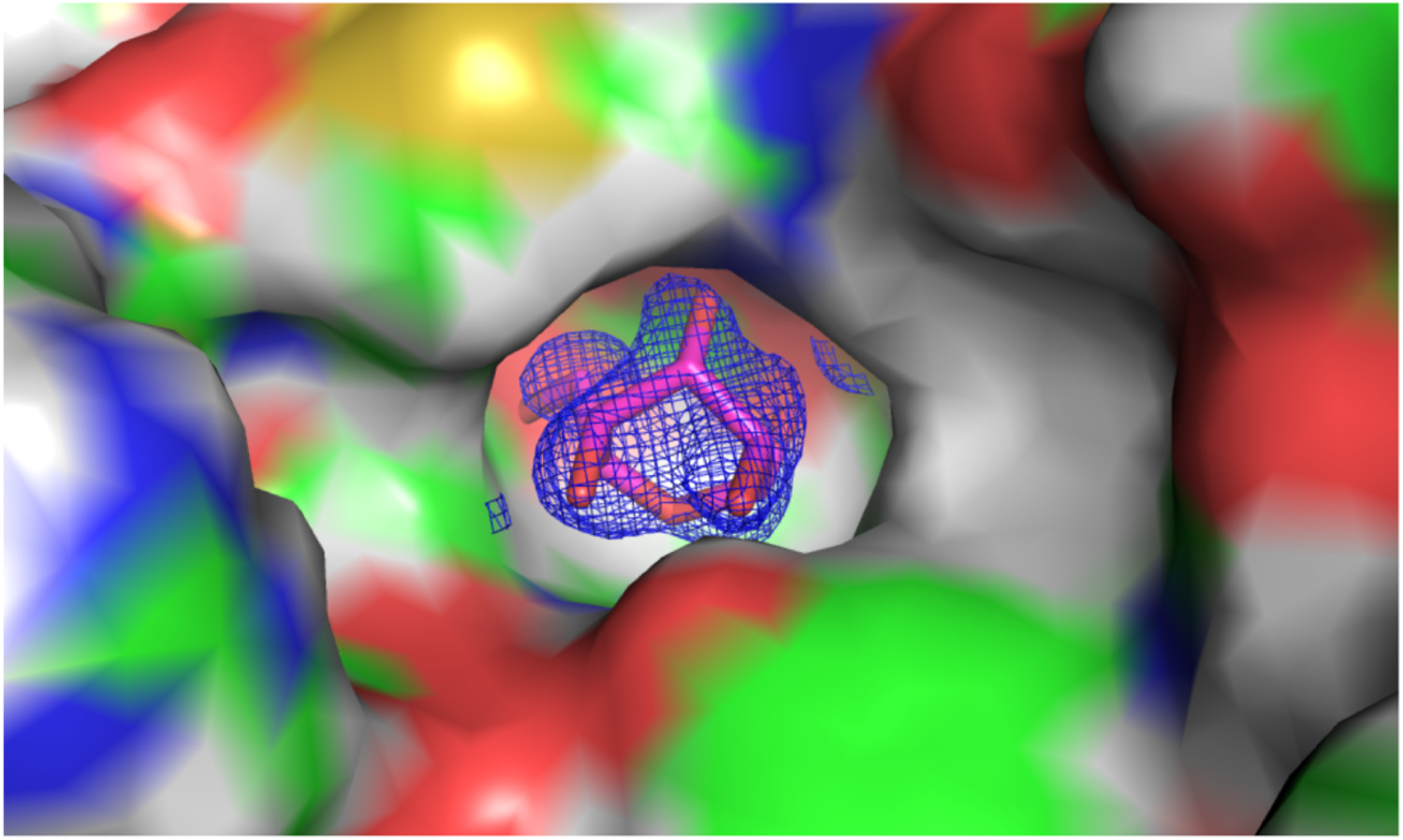
2Fo-Fc map of sugar bound into the S-pocket of Ldt_Mt2_ in the crystal structure. S-pocket is represented in surface. Sugar is shown in stick model in pink color. 2Fo-Fc map is shown in blue color.

**Figure S2:**
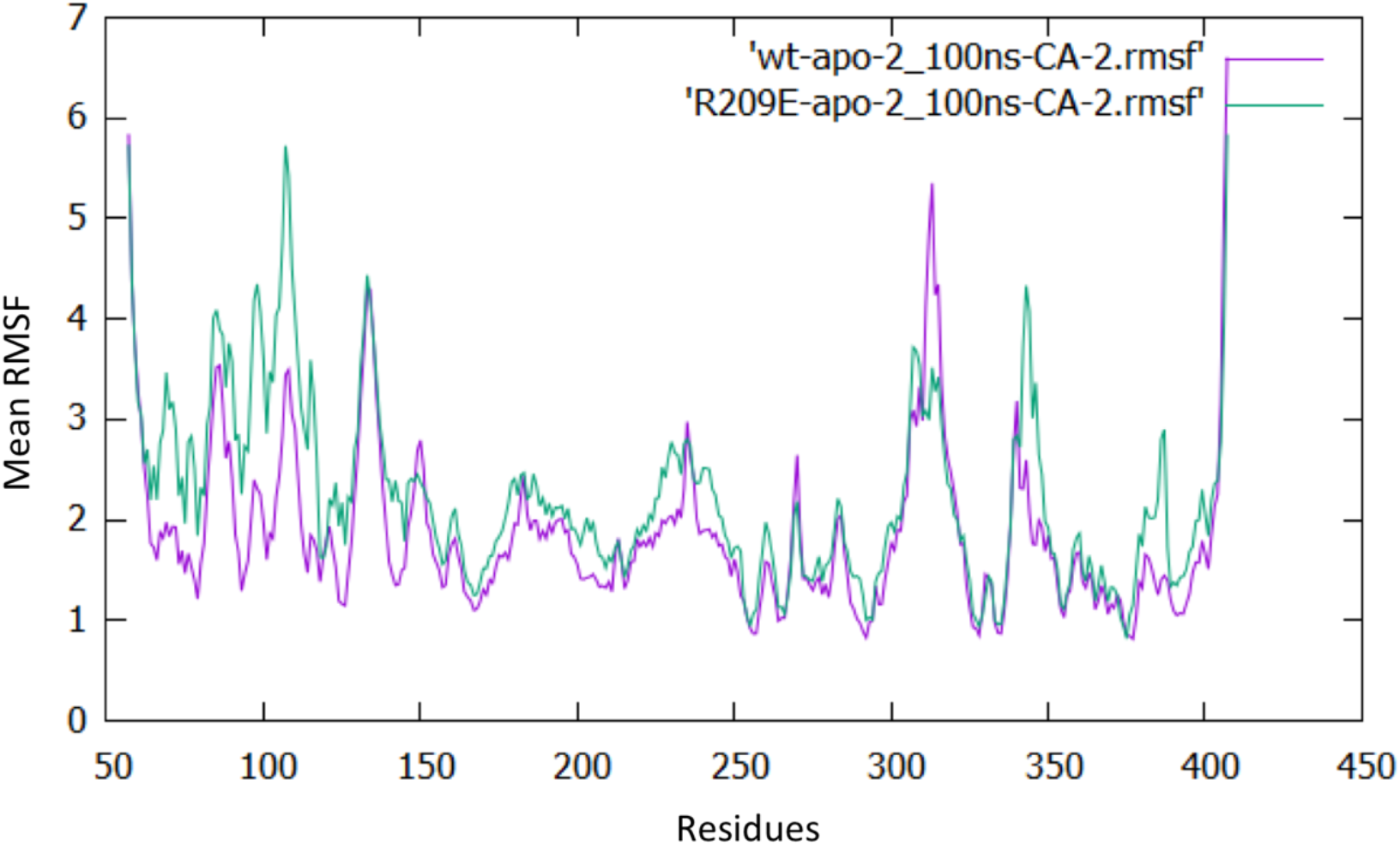
RMSF graph of wild-type Ldt_Mt2_ (purple) and R209 mutant (green) after 100ns of MD simulations.

**Figure S3:**
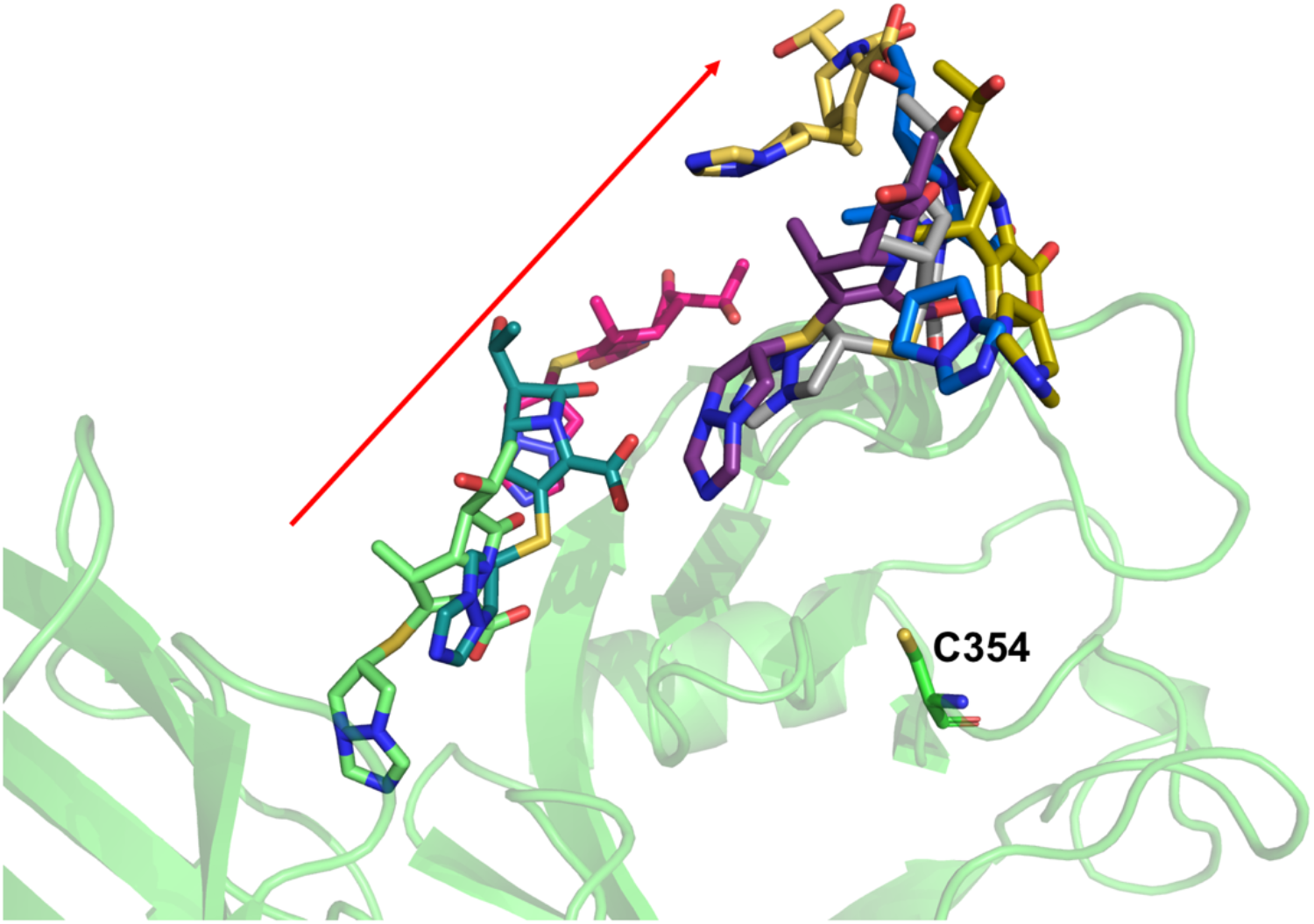
Molecular dynamic trajectory of biapenem bound in S-pocket alone. A red arrow indicates the direction of movement of biapenem during an overall 75 ns of MD simulation. Ldt_Mt2_ is represented in cartoon in green colour and biapenem in various trajectories is represented in stick model.

**Figure S4:**
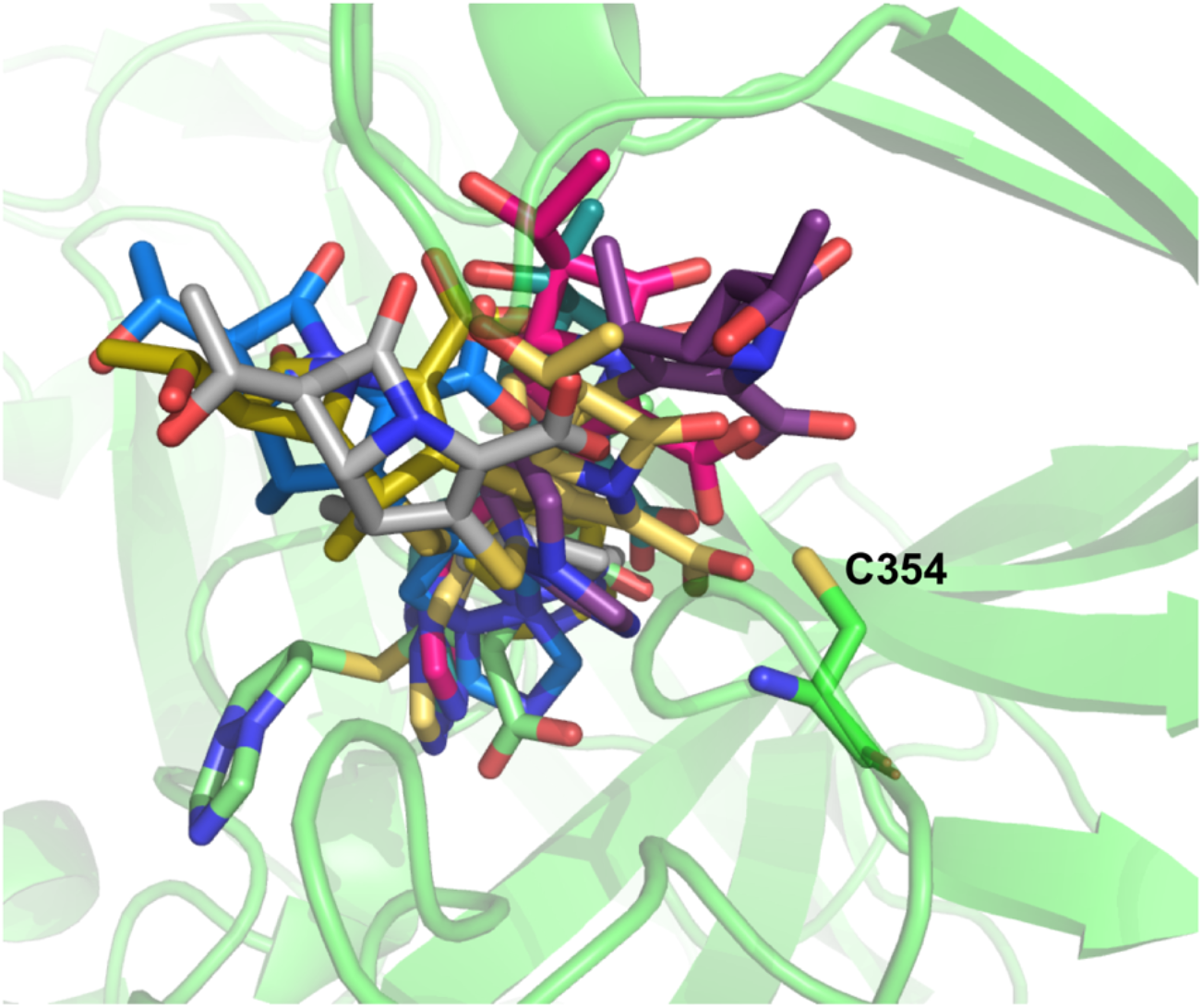
Molecular dynamic trajectory of biapenem bound in catalytic site alone in Ldt_Mt2_. Ldt_Mt2_ is represented in cartoon in green colour and biapenem in various trajectories is represented in stick model.

**Figure S5:**
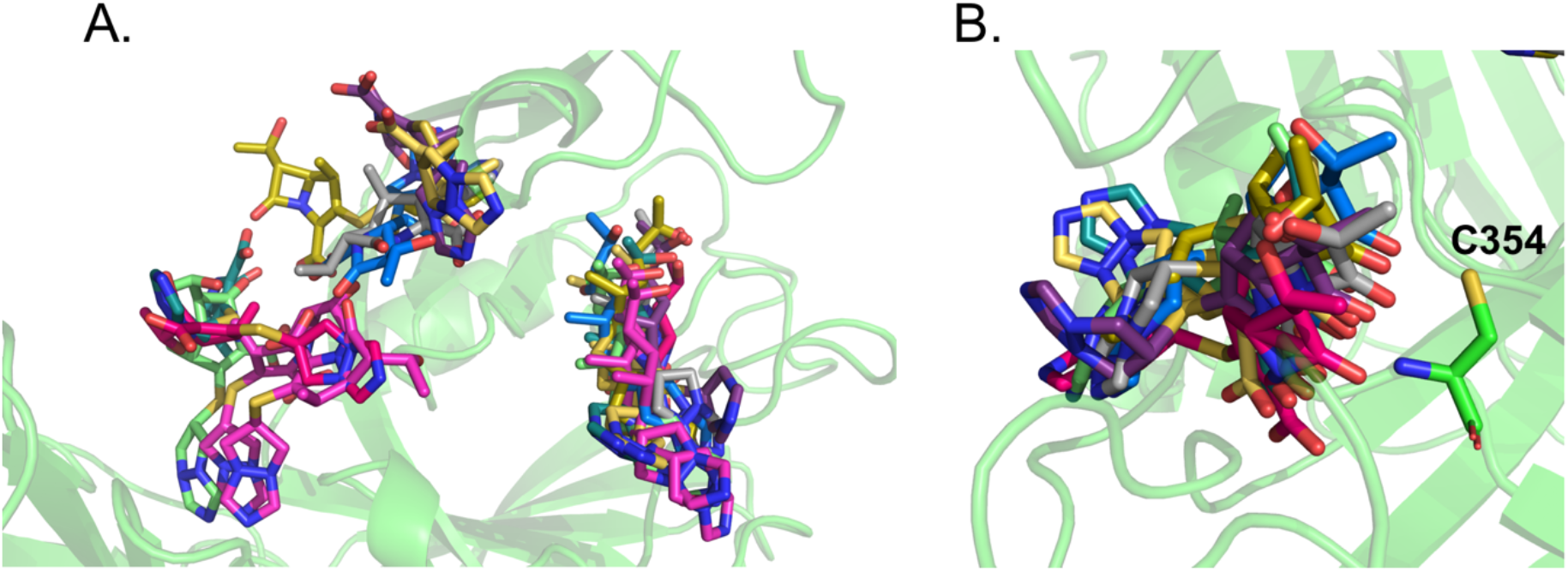
Molecular dynamic trajectory of dual biapenem bound together in both S-pocket and catalytic site. (A) Snapshots of biapenem from both S-pocket and catalytic site (B) Snapshots of biapenem from catalytic site. Ldt_Mt2_ is represented in cartoon in green colour and biapenem in various trajectories is represented in stick model.

**Figure S6:**
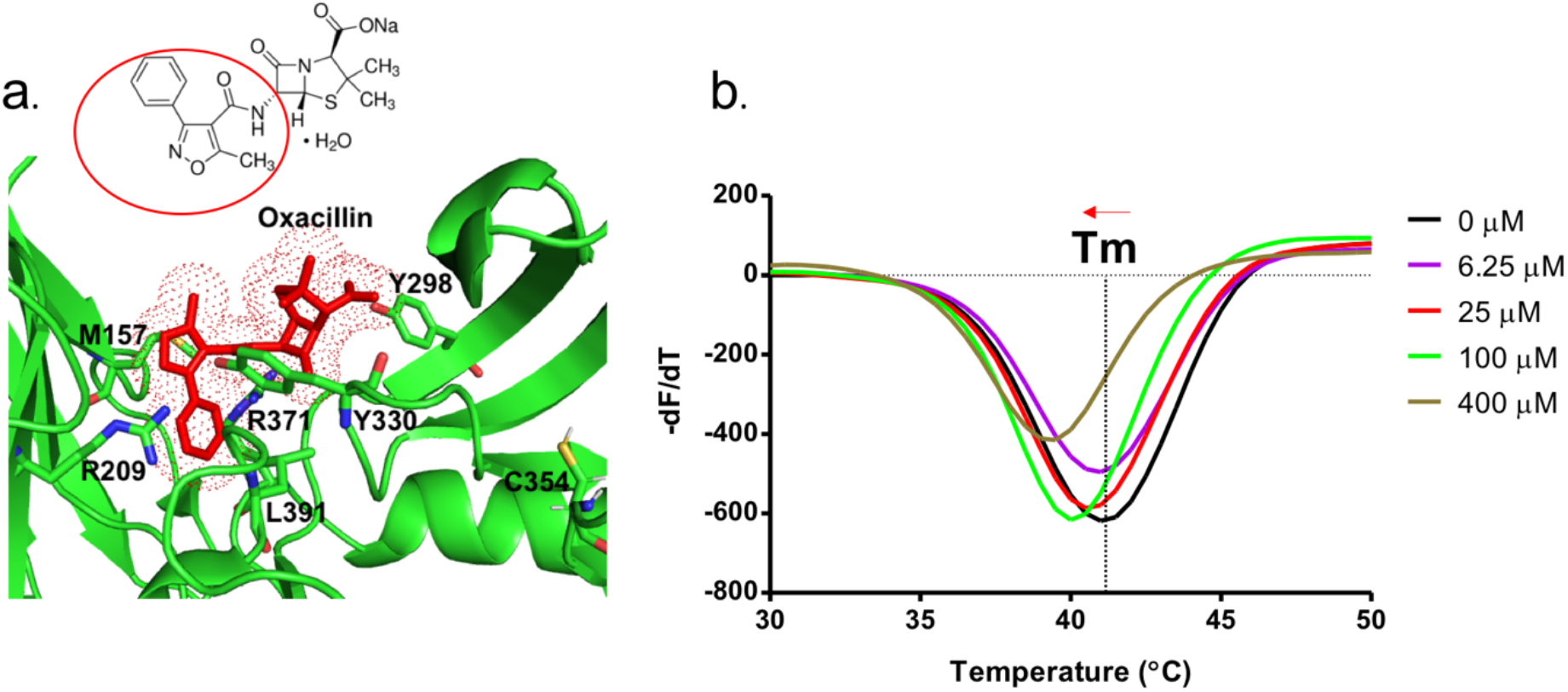
(a) Oxacillin (stick model, red colour) bound into the S-pocket (green colour) of Ldt_Mt2_ through R1 group side chain 5-methyl-3-phenyl-1,2-oxazole-4-carbonyl). 5-methyl-3-phenyl-1,2-oxazole-4-carbonyl group is highlighted with red circle in the chemical structure of oxacillin. Picture in the bottom shows ThermoFluor assays for binding studies of oxacillin with wild-type Ldt_Mt2_.

**Figure S7:**
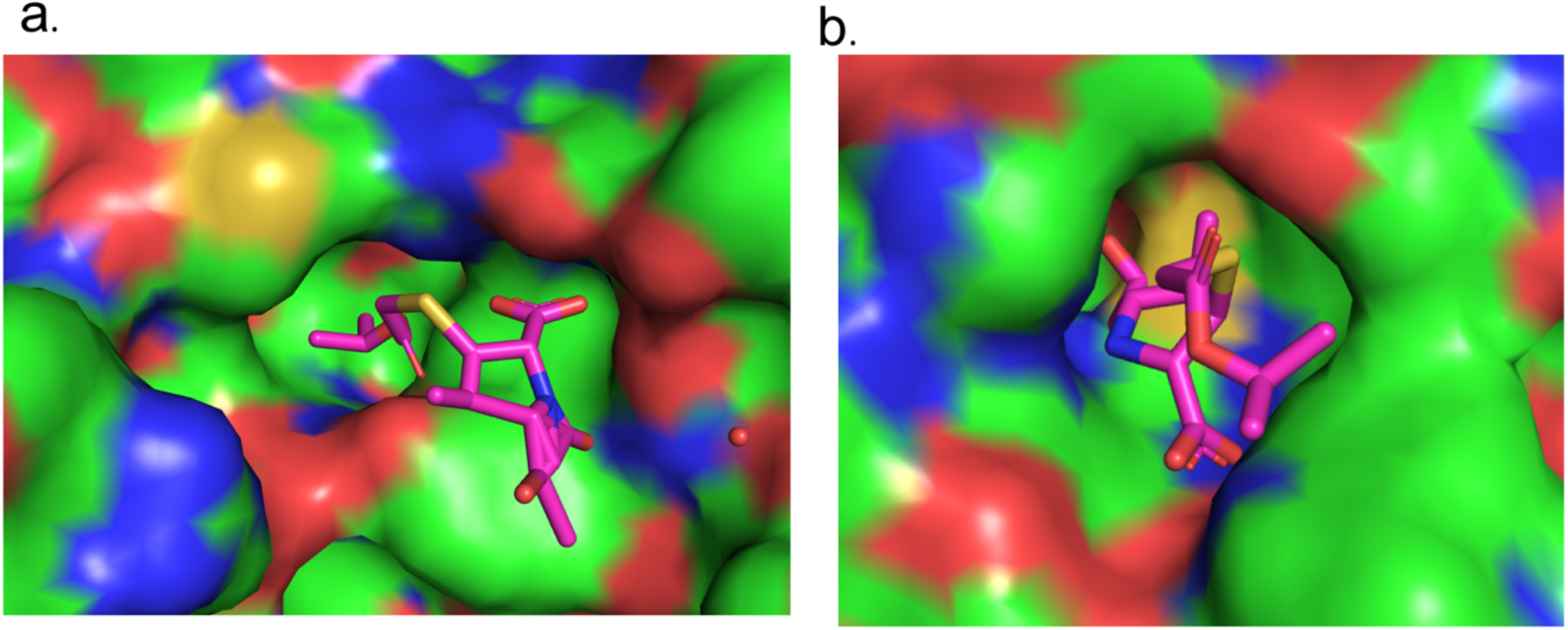
**(A)** T203 drug bound into the S-pocket of Ldt_Mt2_ in the crystal structure. (B) T203 drug bound into the catalytic site of Ldt_Mt2_ in the crystal structure. Ldt_Mt2_ is represented in surface and T203 drug is represented in stock model with pink color.

**Figure S8.**
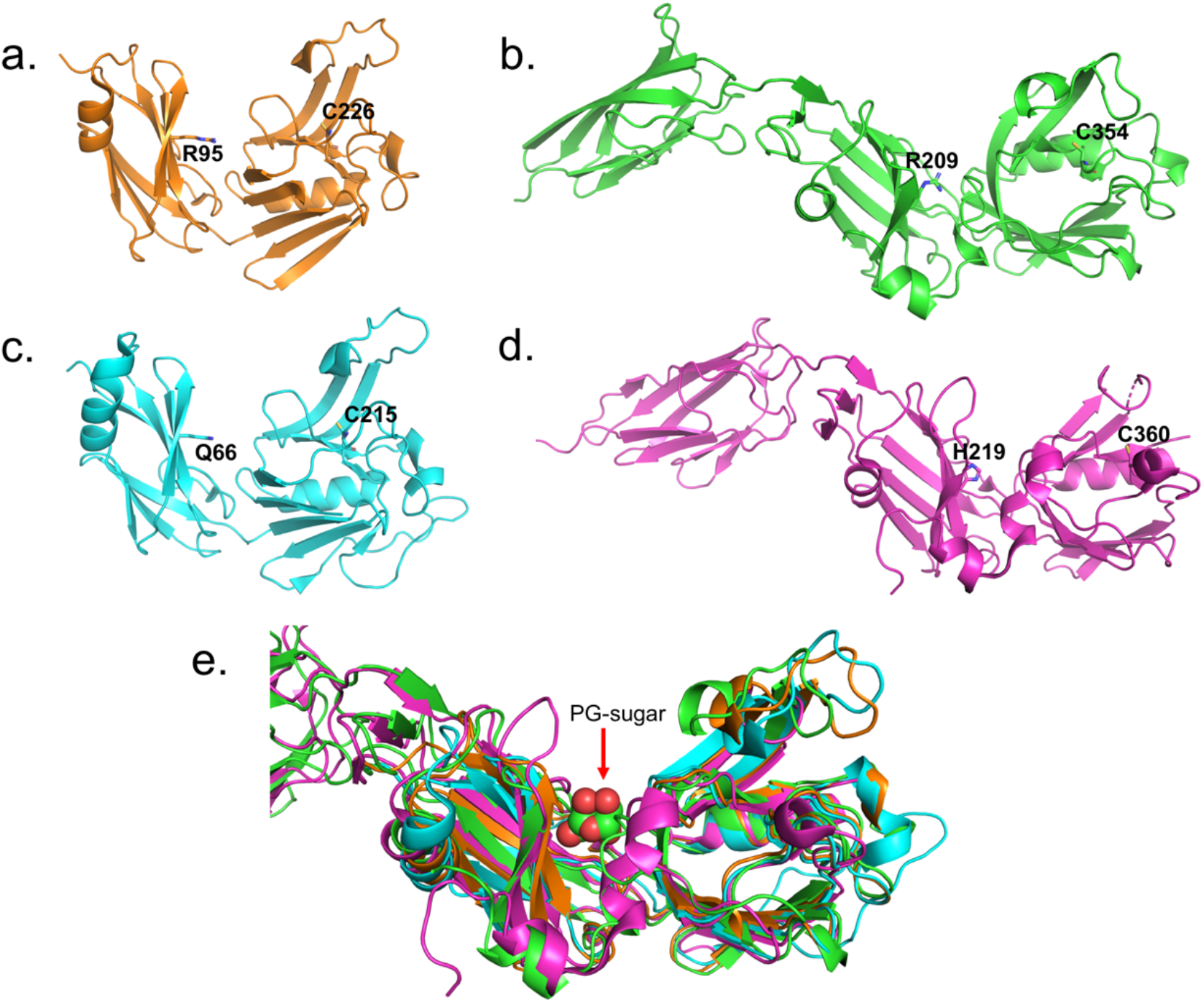
(A-D) Crystal structures of various LDTs, Ldt_Mt1_ (PDB ID: 4JMN; orange color), Ldt_Mt2_ (PDB ID: 7F71; green color), Ldt_Mt3_ (PDB ID: 6D4K; cyan color) and Ldt_Mt5_ (PDB ID: 6D5A; pink color). Highlighted residues in the crystal structures belong to S-pocket and catalytic site. (E) Superposition of various LDTs. PG sugar moiety bound across the S-pocket is shown in sphere model.

## Notes

### Competing Interest Statement

The authors have declared no competing interest.

